# Metabolic adaptations underpin resistance to histone acetyltransferase inhibition

**DOI:** 10.1101/2022.08.12.503669

**Authors:** Timothy R. Bishop, Chitra Subramanian, Eric M. Bilotta, Leopold Garnar-Wortzel, Anissa R. Ramos, Yuxiang Zhang, Joshua N. Asiaban, Christopher J. Ott, Charles O. Rock, Michael A. Erb

## Abstract

Histone acetyltransferases (HAT) catalyze the acylation of lysine side chains and are implicated in diverse human cancers as both oncogenes and non-oncogene dependencies^1^. Acetyl-CoA-competitive HAT inhibitors have garnered attention as potential cancer therapeutics and the first clinical trial for this class is ongoing (NCT04606446). Despite broad enthusiasm for these targets, notably including CBP/p300 and KAT6A/B^2–5^, the potential mechanisms of therapeutic response and evolved drug resistance remain poorly understood. Using comparative transcriptional genomics, we found that the direct gene regulatory consequences of CBP/p300 HAT inhibition are indistinguishable in models of intrinsically hypersensitive and insensitive acute myeloid leukemia (AML). We therefore modelled acquired drug resistance using a forward genetic selection and identified dysregulation of coenzyme A (CoA) metabolism as a facile driver of resistance to HAT inhibitors. Specifically, drug resistance selected for mutations in *PANK3*, a pantothenate kinase that controls the rate limiting step in CoA biosynthesis^6^. These mutations prevent negative feedback inhibition, resulting in drastically elevated concentrations of intracellular acetyl-CoA, which directly outcompetes drug-target engagement. This not only impacts the activity of structurally diverse CBP/p300 HAT inhibitors, but also agents related to an investigational KAT6A/B inhibitor that is currently in Phase-1 clinical trials. We further validated these results using a genome-scale CRISPR/Cas9 loss-of-function genetic modifier screen, which identified additional gene-drug interactions between HAT inhibitors and the CoA biosynthetic pathway. Top hits from the screen included the phosphatase, *PANK4*, which negatively regulates CoA production and therefore suppresses sensitivity to HAT inhibition upon knockout^7^, as well as the pantothenate transporter, *SLC5A6*^8^, which enhances sensitivity. Altogether, this work uncovers CoA plasticity as an unexpected but potentially class-wide liability of anti-cancer HAT inhibitors and will therefore buoy future efforts to optimize the efficacy of this new form of targeted therapy.

---

HATs regulate transcriptional activity by catalyzing the acylation of lysine residues on histone tails and other substrate proteins^1^. Multiple HAT proteins are implicated in the maintenance and emergence of human malignancies and have therefore been targeted for therapeutic development. The highly homologous enhancer factors, CBP and p300, are crucial for the pathogenesis of diverse human cancers, both as non-oncogene dependencies and as part of oncogenic fusion proteins^9–14^. They are also recurrently mutated in lymphomas and lung cancers resulting in a synthetic lethality between the two paralogs^15^. Likewise, several MYST-family HAT proteins, such as KAT6A/B and HBO1, are involved in oncogenic fusions and/or are required for the maintenance of tumorigenic gene expression^10,16,17^. These genetic data have motivated widespread efforts to address HAT proteins with chemical probe discovery and drug development. Multiple structurally diverse chemical probes and clinical-stage inhibitors targeting CBP/p300 and MYST-family HAT domains have now been discovered, supporting translational experiments that further substantiate HATs as anti-cancer drug targets^2–4,18,19^. However, potential biomarkers of response remain as-yet unclear and little is understood about how resistance to these agents may arise in a clinical setting. Here, we employed multiple forward genetic approaches to model the potential mechanisms of resistance to CBP/p300 HAT inhibition. These experiments uncovered dysregulation of CoA biosynthesis as a facile mechanism of resistance to HAT inhibition, revealing a likely universal liability of acetyl-CoA-competitive HAT inhibitors.

## Results

A-485 is a first-in-class chemical probe that inhibits CBP/p300 HAT domains by binding competitively with acetyl-CoA. Large-scale profiling of its anti-proliferative activity in human cancer cell lines has previously demonstrated that hematological malignancies, including multiple myeloma and AML, are among the most sensitive to CBP/p300 HAT inhibition^2^. However, whereas most multiple myeloma cell lines are responsive to A-485, AML includes examples of both hypersensitive and intrinsically resistant cell lines. Due to this heterogeneity, we reasoned that AML would provide an ideal model system to study response and resistance to CBP/p300 HAT inhibition. We first confirmed these data with a panel of AML cell lines **(Fig. 1a)**, demonstrating that sensitive lines exhibited strong G1 cell cycle arrests in response to mid-nanomolar A-485 treatment, whereas resistant lines showed only mild responses at much higher concentrations of the drug **(Extended Data Fig. 1a,b)**. Moreover, while our analyses of publicly available genome-scale CRISPR/Cas9-based essentiality screens^20,21^ indicated that *EP300* is a selective non-oncogene dependency in AML **(Fig. 1b)**, we found that individual AML cell lines vary widely in their sensitivity to *EP300* disruption **(Extended Data Fig. 1c-e)**. Having validated the heterogenous nature of AML responses to genetic and pharmacologic disruption of CBP and/or p300, we next proceeded with comparative transcriptional genomics experiments to understand the gene regulatory impacts of HAT inhibition in sensitive and insensitive AML models.

**Figure 1:**
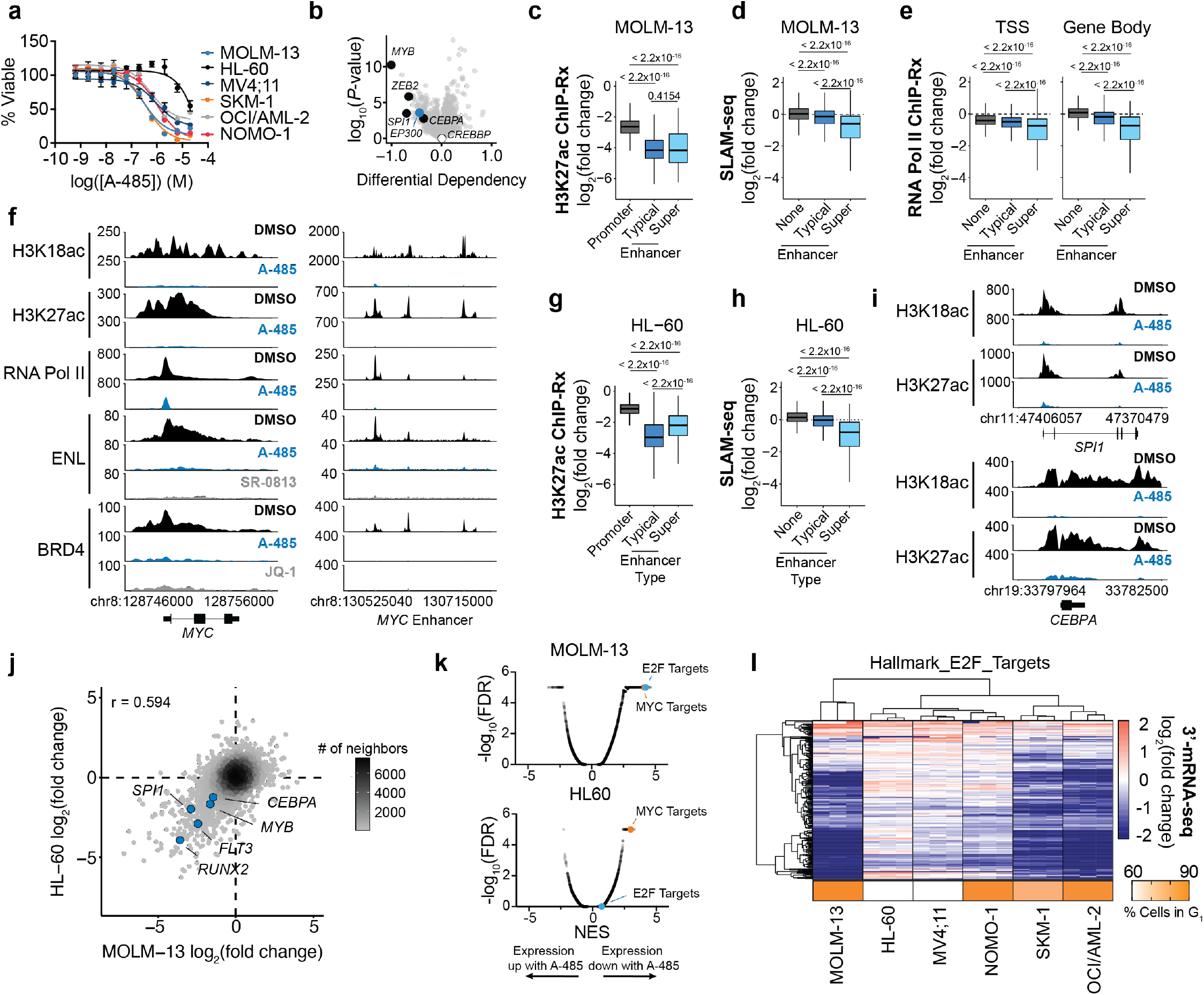
CBP/p300 HAT inhibition disrupts enhancer-driven transcription in sensitive and insensitive AML. **a**, DMSO-normalized cellular viability in a panel of AML cell lines after 72-h treatment with A-485. Mean ± SEM, n = 3. **b**, Volcano plot depicting genome-wide dependency differences between AML cell lines and all other lineages. Data from Cancer Dependency Map release 22Q1. Highlighted in blue are known leukemic driver genes and dependencies. **c**, Boxplots of change in H3K27ac ChIP-Rx signal at promoters (n = 12650), typical enhancers (n = 12823) and super-enhancers (n = 774) after 2-h A-485 (3 μM) treatment in MOLM-13. **d**, Boxplots of DMSO-normalized changes in newly synthesized transcript abundance after 2-h A-485 (3 μM) treatment in MOLM-13 measured by SLAM-seq at genes predicted to be regulated by no enhancer (n = 8151), typical enhancers (n = 3609), and super-enhancers (n = 638). **e**, Boxplots of DMSO-normalized changes in Pol II occupancy at the TSS and in the gene body after 2-h A-485 (3 μM) treatment at genes predicted to be regulated by no enhancer (n = 11028), typical enhancers (n = 3960), and super-enhancers (n = 451). **f**, Gene tracks of ChIP-Rx signal (reference-adjusted reads per million, RRPM) in MOLM-13 at representative loci. **g**, Boxplots of change in H3K27ac ChIP-Rx signal at promoters (n = 12171), typical enhancers (n = 11635) and super-enhancers (n = 681) after 2-h A-485 (3 μM) treatment in HL60.. **h**, Boxplots of DMSO-normalized changes in newly synthesized transcript abundance after 2-h A-485 (3 μM) treatment in HL-60 measured by SLAM-seq at genes predicted to be regulated by no enhancer (n = 7636), typical enhancer (n = 3344), and super-enhancers (n = 592). **i**, Gene tracks of ChIP-Rx signal (RRPM) in HL-60 at representative loci. **j**, Comparison of SLAM-seq responses in MOLM-13 and HL-60 following 2-h of A-485 (3 μM) treatment. **k**, Gene set enrichment analysis (GSEA) of gene-expression changes induced by 48-h A-485 (3 μM) treatment in MOLM-13 and HL-60. **l**, Heatmap representation of DMSO-normalized fold changes in spikein normalized gene expression of E2F target genes and G_1_ cell cycle arrest following 48-h A-485 (3 μM) treatment in a panel of AML cell lines. For all boxplots, boxes represent 25-75 percentiles with whiskers extending to 1.5 * the interquartile range (IQR) and *P* values were determined by a two-tailed Welch’s t-test.

In MOLM-13 cells – a representative example of hypersensitive AML – we found that CBP/p300 HAT inhibition reduced the acetylation of the CBP/p300 substrates, H3K18ac and H3K27ac, globally **(Extended Data Fig. 2a-b)**, but the most pronounced effects occurred at enhancer elements **(Fig. 1c and Extended Data Fig. 2c)**. Despite the widespread hypoacetylation that results from CBP/p300 HAT inhibition^22^, the direct transcriptional effects of A-485 treatment were highly selective. As measured by both SH-linked alkylation for the metabolic sequencing of RNA (SLAM-seq) and RNA Polymerase (Pol) II ChIP-Rx, A-485 preferentially suppressed the transcription of super-enhancer-associated genes **(Fig. 1d,e and Extended Data Fig. 2def)**. This resulted in strong repression of a relatively small subset of genes, notably including several known AML drivers, such as *MYB, FLT3, SPI1, ZEB2*, and *CEBPA*^23^ **(Fig. 1j and Extended Data Fig. 2e-f)**. Loss of Pol II signal at these genes was observed at both the transcription start site (TSS) and throughout the gene body, suggesting that rather than a defect in Pol II pause release, CBP/p300 HAT inhibition results in a loss of Pol II recruitment to the promoters of enhancer-regulated genes **(Extended Data Fig. 2g)**. Despite similar levels of histone hypoacetylation at typical and super enhancers, the transcription of super-enhancer regulated genes was more effectively suppressed by CBP/p300 HAT inhibition, suggesting that their regulation is more sensitive to CBP/p300-driven acetylation events.

To understand the mechanisms underlying the reduced activity of RNA Pol II, we performed ChIP-Rx for ENL and BRD4, chromatin reader proteins that bind acetyl-lysine side chains and are implicated in leukemia pathogenesis^24–27^. After 2 hours of A-485 treatment, the occupancy of ENL and BRD4 on chromatin was severely diminished, suggesting that higher-order transcriptional machinery is compromised **(Fig. 1f and Extended Data Fig. 2h-k)**. Consistent with the greater loss of histone acetylation from enhancers, BRD4 was more strongly displaced from enhancers than promoters. Furthermore, displacement of BRD4 occurred disproportionately at super-enhancer elements compared to typical enhancers **(Extended Data Fig. 2j)**. Since BRD4 is required for RNA Pol II activity globally^28,29^, selectively disrupting its localization to enhancers by inhibiting CBP/p300 HAT domains might explain, at least in part, the selective suppression of lineage-specifying transcripts controlled by super-enhancers. The chromatin localization of ENL, which recruits the super elongation complex to the promoters of key leukemogenic drivers^24^, was also disrupted by CBP/p300 HAT inhibition globally, but with pronounced effects at promoters where ENL binds most abundantly **(Extended Data Fig. 2k)**. In all, these data suggest that CBP/p300 HAT inhibition and concomitant histone hypoacetylation lead to eviction of acetyl-lysine binding proteins, deficient Pol II recruitment and activity, and a loss of enhancer-mediated leukemogenic transcription. These data are broadly consistent with results from other cell types^18,30,31^, but it remains unclear whether the changes we and others observe are sufficient to explain the anti-proliferative effects of CBP/p300 HAT inhibition. Comparison to lineage-matched insensitive controls have not been described and, therefore, the potential for these phenotypes to act as predictive biomarkers of response remains untested.

We hypothesized that the acute effects of CBP/p300 HAT inhibition would be less severe in cell lines that are insensitive to the antiproliferative effects of A-485 treatment. However, while proliferation in the presence of A-485 varies widely across AML cells lines, there was little difference in the global depletion of H3K27ac observed by immunoblot upon CBP/p300 HAT inhibition **(Extended Data Fig. 3a,b)**. Like we observed in hypersensitive MOLM-13 cells, ChIP-Rx of H3K18ac and H3K27ac in the A-485-tolerant AML cell line, HL-60, revealed a global depletion of H3K18ac and H3K27ac, with pronounced effects at enhancers compared to promoters **(Fig. 1g and Extended Data Fig 3c,d)**. Moreover, the transcriptional effects of CBP/p300 HAT inhibition detected by SLAM-seq in HL-60 cells were indistinguishable from MOLM-13 cells, in both the identity of the suppressed transcripts as well as their overall magnitude of change **(Fig. 1h and Extended Data Fig. 3e,f)**. A comparable number of genes were significantly downregulated by A-485 treatment and the overall changes were highly correlated between the two cell lines **(Fig. 1j and Extended Data Fig. 3e,f)**. As in MOLM-13 cells, many of the most severely down-regulated transcripts in HL-60 cells were known AML drivers and there was a strong preference for disruption of super-enhancer mediated transcription **(Fig. 1h)**. Our results revealed identical acute responses to HAT inhibition, but to determine if a divergent transcriptional response would become apparent at steadystate, we performed 3’-end mRNA sequencing following 48 hours of A-485 exposure in a panel of 6 AML cell lines. These data detected highly divergent effects in sensitive and resistant cell lines. For example, gene set enrichment analysis revealed an enrichment for down-regulation of MYC target genes in all cell lines, but only sensitive cell lines that underwent a strong G1 cell cycle arrest also repressed E2F transcriptional targets **(Fig. 1k,l and Extended Data Fig. 4a,b)**. These data indicate that the primary transcriptional response to A-485 treatment, including the selective suppression of leukemia proto-oncogenes, is not predictive of, or sufficient for, an anti-proliferative phenotypic response. The decoupling of the primary transcriptional response from compound sensitivity indicates that lineage and/or state-dependent adaptations occurring secondary to the initial transcriptional insult ultimately determine the sensitivity of cells to HAT inhibition, similar to previous observations for BET bromodomain inhibition^32^.

To provide an unbiased assessment of the mechanisms that determine response and resistance to HAT inhibition, we next modeled acquired resistance to A-485 using a forward genetic selection. Specifically, we cultured MOLM-13 cells in the presence of A-485, incrementally increasing the concentration of drug from 50 nM to 2 μM (hereafter referred to as MOLM-13-2R cells) while simultaneously growing a control population treated with DMSO to the same high passage number (MOLM-13-HP). While A-485 inhibited the viability of the parental and MOLM-13-HP cell lines nearly identically (half-maximal inhibitory concentrations (IC_50_) of 310 nM and 476 nM, respectively), MOLM-13-2R displayed a 70-fold shift in sensitivity (IC_50_ = 23.1 μM) **(Fig. 2a)**. Furthermore, extended treatment with A-485 over 2 weeks revealed that MOLM-13-2R cells can proliferate exponentially in concentrations of at least 3 μM, which was cytotoxic to MOLM-13-HP cells **(Fig. 2b)**. We observed cross-resistance to the chemically distinct CBP/p300 HAT inhibitor CPI-1612^33^ **(Fig. 2a)** and no change in resistance to A-485 by co-treatment with the MDR1 inhibitor, verapamil, supporting the conclusion that resistance is not driven by drug efflux **(Fig. 2c)**. However, we found that MOLM-13-2R cells remained dependent on CBP/p300 function as they were equivalently sensitive to the CBP/p300 bromodomain inhibitors, GNE-049, GNE-781, GNE-272 and SGC-CBP30^34–37^, as well as a CBP/p300 degrader, dCBP-1^3^, which is derived from a CBP/p300 bromodomain ligand **(Fig. 2d,e)**. Furthermore, in a screen of 472-compounds targeting chromatin and transcription regulatory proteins, we detected no other changes in sensitivity **(Fig. 2e)**. Altogether, these data established that resistance was specific to HAT inhibitors and motivated the hypothesis that changes in the MOLM-13-2R cell line were affecting drug-target engagement.

**Figure 2:**
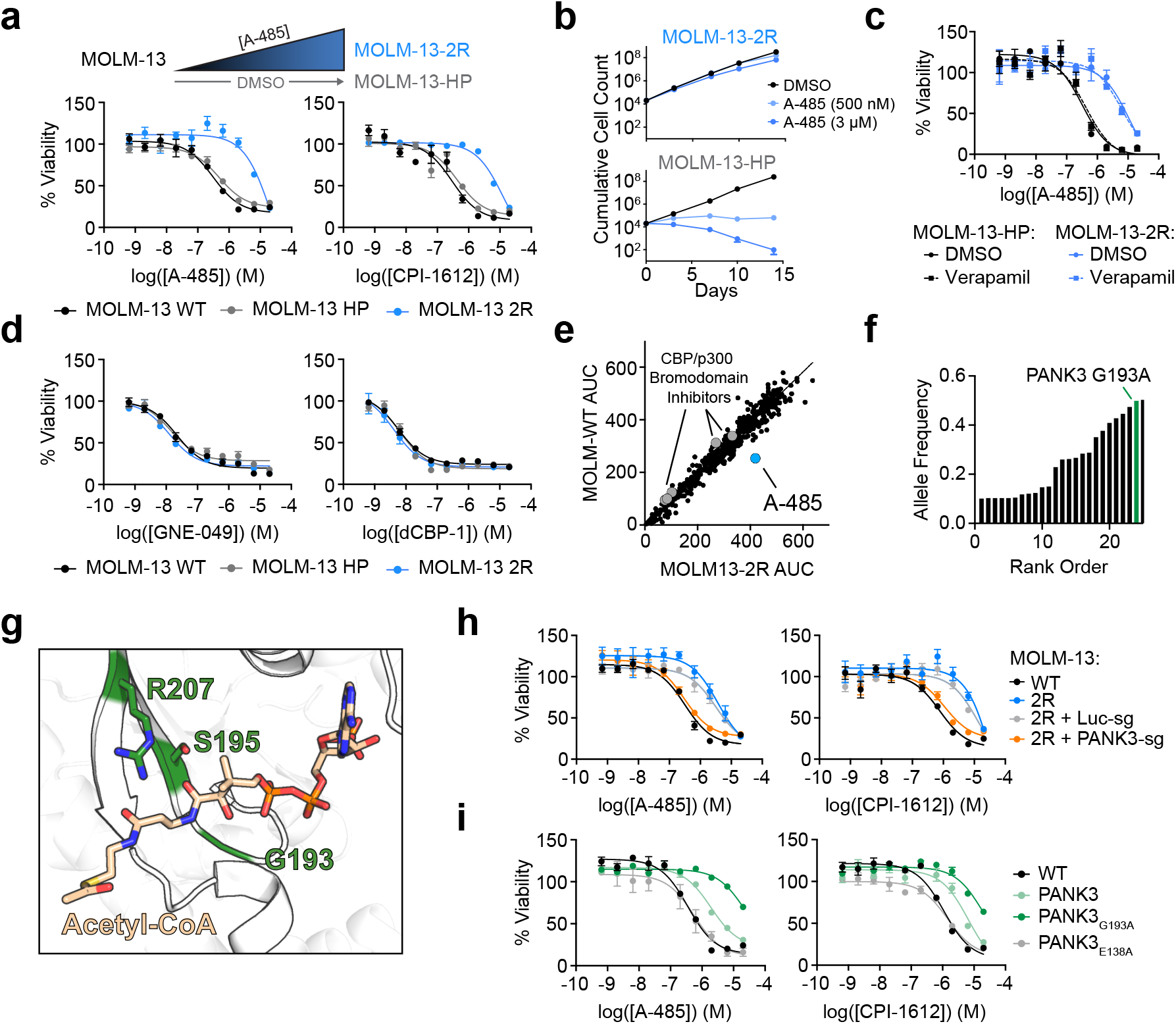
*PANK3* alterations are necessary and sufficient for acquired resistance to HAT inhibition. **a,** DMSO-normalized cellular viability in MOLM-13, MOLM-13-2R, and MOLM-13-HP after 72-h treatment with indicated compounds as determined by ATP-lite assay. Mean ± SEM, n = 3. **b**, Proliferation of MOLM-13-2R and MOLM-13-HP in response to A-485 treatment over 14 days. **c**, DMSO-normalized cellular viability of MOLM-13-2R and MOLM-13-HP with A-485 and verapamil (10 μM) co-treatment as determined by ATP-lite assay. Mean ± SEM, n = 3. **d**, DMSO-normalized cellular viability in MOLM-13, MOLM-13-2R, and MOLM-13-HP after 72-h treatment with indicated compounds as determined by ATP-lite assay. Mean ± SEM, n = 3. **e**, Sensitivity of MOLM-13 and MOLM-13-2R cells to 72-h treatment with compounds from epigenetics-targeting library. Values represent area under curve from 10-point dose responses. **f**, Rank ordered allele frequencies of variants called by whole-exome sequencing with alteration predicted to lead to PANK3 G193A (chr5 167993075 C>G) mutation highlighted. **g**, Structural view of the acetyl-CoA and pantothenate binding site in PANK3 (PDB: 3MK6). Amino acids G193, S195V, and R207 are highlighted in green. **h**, DMSO-normalized cellular viability in MOLM-13, MOLM-13-2R, and MOLM-13-2R sgRNA-expressing populations after 72-h treatment with A-485 and CPI-1612 as determined by ATP-lite assay. Mean ± SEM, n = 3. **i**, DMSO-normalized cellular viability in MOLM-13 expressing PANK3 ORFs after 72-h treatment with A-485 and CPI-1612 as determined by ATP-lite assay. Mean ± SEM, n = 3.

To identify the alleles potentially responsible for this resistance, we performed whole-exome sequencing (WES) in the parental and resistant cell lines **(Fig. 2f and Extended Data Fig. 5a,b)**. Because MOLM-13-2R still require CBP/p300 function for survival, we hypothesized that we would identify alterations within CBP and/or p300 that prevent drug-target engagement. Indeed, we discovered an S1650F mutation in the HAT domain of p300 **(Extended Data Fig. 5b)**, but it occurred at a low estimated allele frequency (~10%) and CRISPR-base editing invalidated it as a driver of resistance to HAT inhibition **(Extended Data Fig. 5c,d)**. Prioritizing high-frequency alleles for further validation, we detected a G193A mutation in pantothenate kinase 3 (PANK3), which occurred at an estimated allele frequency of 49.9% **(Fig. 2f)**. This enzyme, along with its paralogs, PANK1 and PANK2, catalyzes the first and rate limiting step in CoA biosynthesis by converting pantothenate (vitamin B5) to 4’-phosphopantothenate^6,38^. Orthosteric feedback inhibition of PANK enzymes by acetyl-CoA is a major determinant of the intracellular CoA concentration^38,39^, and the G193A mutation in PANK3 is located at the site of both pantothenate and acetyl-CoA binding **(Fig. 2g and Extended Data Fig. 6a)**. Therefore, we hypothesized that this *PANK3* mutation might affect the cellular production of acetyl-CoA, the CBP/p300 substrate that binds competitively with A-485. Using CRISPR/Cas9 to knockout *PANK3* from MOLM-13-2R cells, we confirmed that *PANK3_G193A_* is necessary for resistance to A-485 and CPI-1612. **(Fig. 2h and Extended Data Fig. 6b)**. To test if the PANK3_G193A_ mutation was sufficient to drive resistance to CBP/p300 HAT inhibitors, we stably expressed wild-type *PANK3, PANK3_G193A_*, and a catalytically inactive *PANK3* mutant (PANK3_E138A_)^40^ in parental MOLM-13 cells by lentiviral transduction **(Extended Data Fig. 6c)**. While cell lines expressing PANK3_E183A_ remained as sensitive to HAT inhibition as the parental cell line, overexpression of PANK3_WT_ and PANK3_G193A_ drove robust resistance to HAT inhibition, with 5- and 50-fold decreases in sensitivity to A-485, respectively **(Fig. 2i)**. Overexpressing PANK3_WT_ and PANK3_G193A_ was also able to confer resistance to CBP/p300 HAT inhibitors in another AML cell line, OCI/AML-2, and the multiple myeloma cell line, RPMI-8226 **(Extended Data Fig. 6e-h)**. As with MOLM-13-2R cells, isolated expression of PANK3_G193A_ did not alter sensitivity to the CBP/p300 bromodomain inhibitor, GNE-049, or the CBP/p300 degrader, dCBP-1 **(Extended Data Fig. 6d-h)**. These results confirmed that the G193A alteration is necessary and sufficient to confer resistance to CBP/p300 HAT inhibitors and demonstrated that simply overexpressing PANK3 is sufficient to diminish the effects of HAT inhibition.

In parallel, we performed an unbiased CRISPR/Cas9-based modifier screen to identify gene-drug interactions at scale **(Fig. 3a and Extended Data Fig. 7a,b)**. This screen uncovered additional regulators of CoA biosynthesis capable of modulating sensitivity to CBP/p300 HAT inhibitors. Among the top hits that conferred resistance to A-485 treatment was *PANK4*, a pseudo pantothenate kinase that contains a DUF89 family phosphatase domain^41,42^ **(Fig. 3a,b and Extended Data Fig. 7d)**. PANK4 dephosphorylates 4’-phosphopantetheine, an intermediate metabolite of CoA biosynthesis, to antagonize CoA biosynthesis^7^. We confirmed that genetic disruption of *PANK4* conferred a competitive growth advantage under selective pressure from CBP/p300 HAT inhibition and clonal expansions of the MOLM-13 cells with *PANK4* knockouts displayed diminished sensitivity to CBP/p300 HAT inhibition by A-485 **(Fig. 3c,d and Extended Data Fig. 7f,h)**. Furthermore, our analysis of a recently published CRISPR screen in multiple myeloma cells, which used a different sgRNA library than our screen, revealed *PANK4* was a top hit for A-485 resistance in this context as well **(Extended Data Fig. 7c)**^18^.We also discovered that genetic depletion of *SLC5A6*, which encodes the solute carrier responsible for cellular import of pantothenate^8^, enhanced the sensitivity of MOLM-13 cells to A-485 **(Fig. 3a,b and Extended Data Fig. 7e)**. Competitive growth assays confirmed the enhanced sensitivity of MOLM-13 cells to CBP/p300 HAT inhibition following *SLC5A6* disruption by CRISPR/Cas9 **(Fig. 3e)**. Moreover, clonally expanded MOLM-13 cell lines with knockout of *SLC5A6* exhibited enhanced sensitivity to CBP/p300 HAT inhibition **(Fig. 3f and Extended Data Fig. 7g,i)**. Again, *PANK4* and *SLC5A6* knockout did not affect sensitivity to GNE-049 or dCBP-1 **(Extended Data Fig. 7h,i)**. Together, these results further validate the CoA biosynthetic pathway as a primary modulator of sensitivity to CBP/p300 HAT inhibitors **(Fig. 3g)**.

**Figure 3:**
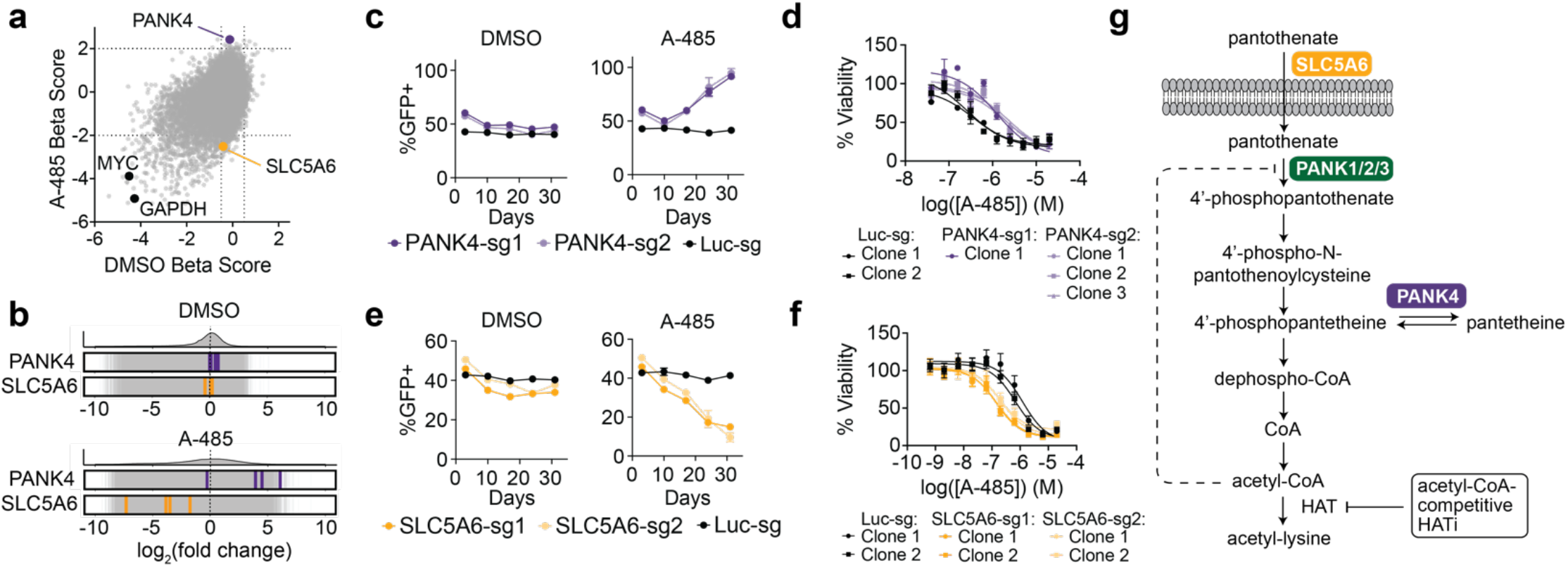
Forward genetic screen converges on gene-drug interactions with the CoA biosynthetic pathway. **a**, Gene beta-scores in A-485 versus DMSO treated populations derived from MAGeCK-mle analysis of pooled CRISPR-Cas9 screen. **b**, Quantification of the change in PANK4- and SLC5A6-targeting sgRNA abundance in A-485 and DMSO treated populations at day 21 compared to day 0. **c**, CRISPR-Cas9 competitive growth assays where proportion of GFP^+^ (PANK4 sgRNA^+^) cells is monitored over time in presence of A-485 or vehicle. Mean ± SEM, n = 3. **d**, DMSO-normalized cellular viability in clonally selected MOLM-13 (Cas9^+^) cells expressing luciferase or PANK4 targeting sgRNAs after 72-h treatment with A-485 as determined by ATP-lite assay. Mean ± SEM, n = 3. **e**, As in **c**, but with SLC5A6 targeting sgRNAs. **f**, As in **d**, but with SLC5A6 targeting sgRNAs. **k**, Schematic depiction of CoA biosynthesis pathway.

We hypothesized that these alterations might affect sensitivity to CBP/p300 HAT inhibitors by modulating intracellular concentrations of acetyl-CoA. Since orthosteric HAT inhibitors bind competitively with acetyl-CoA, its abundance would be predicted to affect drug-target engagement in cells. Targeted metabolomic measurements revealed that MOLM-13-2R, which are heterozygous for *PANK3_G193A_*, contain elevated levels of both CoA and acetyl-CoA **(Fig. 4a and Extended Data Fig. 8a)**. In fact, exogeneous expression of PANK3 and PANK3_G193A_ was sufficient to increase the production of CoA and acetyl-CoA in MOLM-13 and HEK293T cells **(Fig. 4a and Extended Data Fig. 8a-c)**. Additionally, we detected increases in several other acyl-CoA species including butyryl-CoA, which has recently been shown to augment CBP/p300 activity^43^ **(Extended Data Fig. 8b,c)**. As a surrogate measure of target engagement, we detected bulk H3K27ac abundance by immunoblot following treatment with A-485. While we observed concentration-dependent loss of H3K27ac in parental MOLM-13 cells, PANK3_G193A_ over-expressing MOLM-13 showed a large shift in response requiring much higher concentrations of A-485 to reduce H3K27ac abundance **(Fig. 4b)**. In line with these findings, MOLM-13-2R exhibited a muted transcriptional response to HAT inhibition compared to the parental cell line, with few transcripts changing significantly in response to treatment **(Fig. 4c)**. These results support a model whereby the hypermorphic PANK3_G193A_ mutation drives the accumulation of supraphysiological acetyl-CoA concentrations that effectively compete with the drug at the HAT active site. To validate this orthogonally, we used a smallmolecule PANK agonist, PZ-3022, that increases cellular levels of acetyl-CoA^44^. As expected, treatment with this compound reduced sensitivity to CBP/p300 HAT inhibitors but did not affect sensitivity to CBP/p300 bromodomain inhibitors **(Fig. 4d and Extended Data Fig. 9a)**.

**Figure 4:**
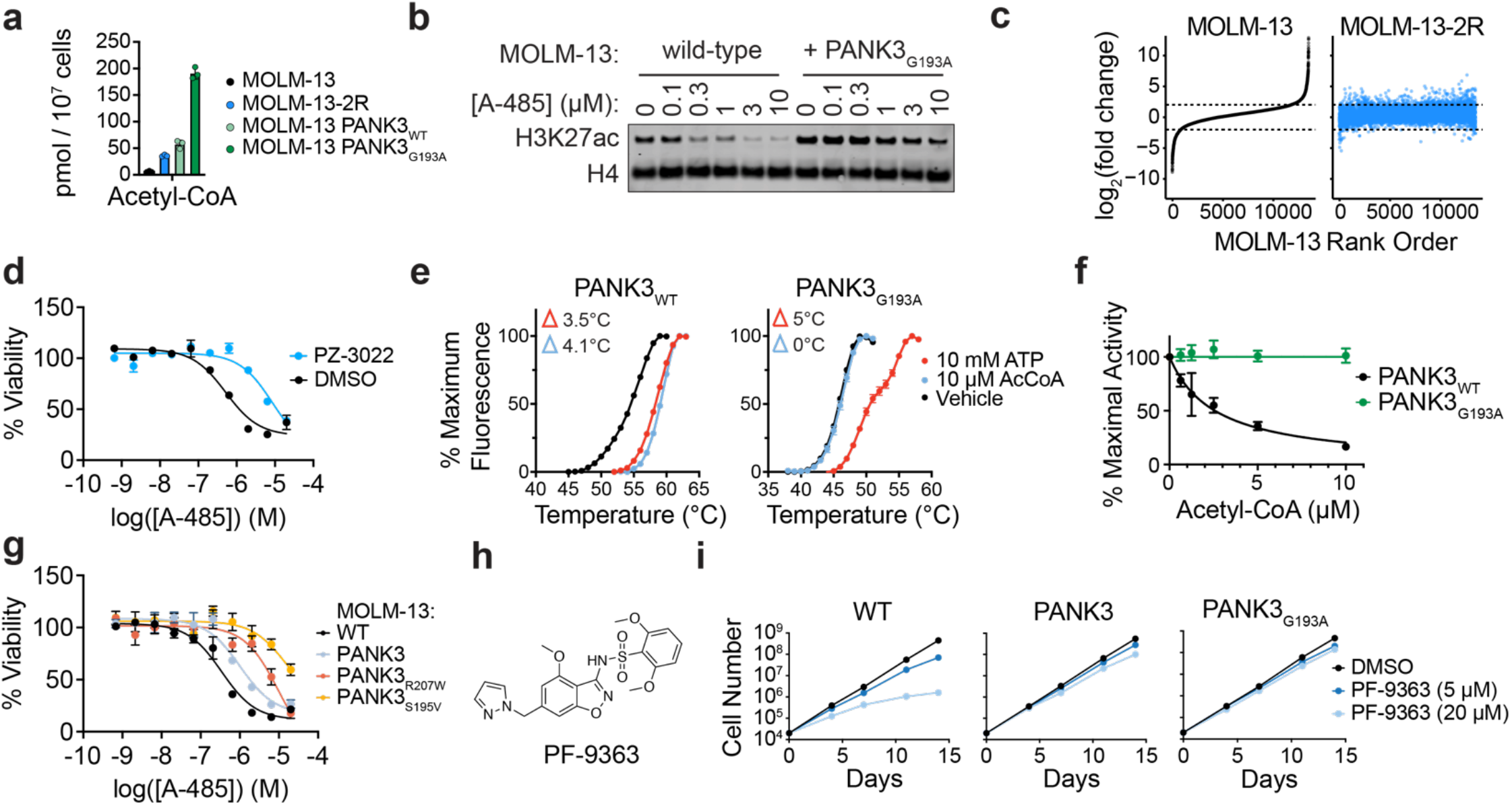
PANK3_G193A_ is a hypermorphic mutant that confers resistance to acetyl-CoA-competitive HAT inhibitors. **a**, Total cellular acetyl-CoA levels determined by mass-spectrometry. Mean ± SEM, n = 3. **b**, Immunoblot for CBP/p300 substrate H3K27ac following 2-h of A-485 treatment. **c**, Expression changes for all genes expressed in either MOLM-13 or MOLM-13-2R (13,506 genes) following 48-h of A-485 (3 μM) treatment. Genes are plotted in ascending order based on fold-change in MOLM-13. **d**, DMSO-normalized cellular viability in MOLM-13 co-treated with A-485 and either PZ-3022 (1 μM) or DMSO for 72-h. Mean ± SEM, n = 3. **e**, Thermal stabilization of PANK3 and PANK3_G193A_ by ATP and acetyl-CoA. **f**, PANK3 and PANK3_G193A_ activity in the presence of acetyl-CoA. Mean ± SEM, n = 3. **g**, DMSO-normalized cellular viability in MOLM-13 expressing PANK3 ORFs after 72-h A-485 treatment. Mean ± SEM, n = 3. **h**, Structure of PF-9363. **i**, Proliferation of MOLM-13 expressing PANK3 ORFs over 14-d of PF-9363 treatment. Mean ± SEM, n = 3.

We next sought to understand mechanistically how the G193A mutation led to this gain of function phenotype. PANK3 activity is normally controlled by negative feedback inhibition, whereby acetyl-CoA occupies the catalytic site that is bound by pantothenate and ATP^38^. Modeling the G193A mutation onto the PANK3 crystal structure predicted a steric clash that would interfere with both acetyl-CoA and pantothenate binding **(Extended Data Fig. 8d,e)**. Given this, we speculated that PANK3_G193A_ was potentially refractory to feedback inhibition by acetyl-CoA. Differential scanning fluorimetry experiments revealed that while both PANK3_WT_ and PANK3_G193A_ were bound and stabilized by ATP, PANK3_G193A_ was refractory to acetyl-CoA binding **(Fig. 4e)**. Similarly, in radiometric PANK activity assays, acetyl-CoA elicited dose-dependent inhibition of PANK3_WT_ but not PANK3_G193A_ **(Fig. 4f)**. Although PANK3_G193A_ binding to pantothenate is compromised, ATP affinity is increased, preserving catalytic activity (specific activity 5.1 vs 277.1 pmol/min/μg) **(Extended Data Fig. 8f,g)**. Like G193A, other mutations nearby the acetyl-CoA binding tunnel of PANK3, like R207W and S195V, are known to diminish negative feedback inhibition by acetyl-CoA, leading to hyperactive production of CoA and its esters^39,41^. We found that PANK3_S195V_ and PANK3_R207W_ expression also elicited resistance to CBP/p300 HAT inhibition in MOLM-13 cells **(Fig. 4g and Extended Data Fig. 9b)**. In all, these results support a model whereby, despite being less active than PANK3_WT_, PANK3_G193A_ and other PANK3 mutants that are refractory to acetyl-CoA can drive the production of supraphysiological concentrations of acyl-CoA species.

We predicted that modulation of intracellular acetyl-CoA concentrations would be a general mechanism of resistance to all acetyl-CoA-competitive HAT inhibitors. PF-07248144, an inhibitor of the structurally distinct MYST HAT domains of KAT6A/B, is currently being evaluated in Phase I clinical trials for treatment-refractory breast cancer (NCT04606446). *KAT6A* has also been described as a dependency in AML^16^, so we determined if modulating acetyl-CoA levels could shift sensitivity to MYST inhibition in AML cell lines. Indeed, overexpressing PANK3 and PANK3_G193A_ in MOLM-13 cells conferred resistance to the KAT6A/B inhibitor, PF-9363^45^ **(Fig. 4h,i and Extended Data Fig. 9c)**. Resistance to PF-9363 was also observed in the G193A heterozygous MOLM-13-2R cell line and could be ablated by genetic knockout of *PANK3* in this cell line **(Extended Data Fig. 9d)**. Similar to CBP/p300 HAT inhibitors, we also observed a shift in KAT6A/B inhibition by PF-9363 by measuring the abundance of its substrate, H3K23ac, in response to drug treatment **(Extended Data Fig. 9e)**. These results indicate that the facile manipulation of CoA metabolism is relevant not just to CBP/p300 HAT inhibitors, but all acetyl-CoA competitive HAT inhibitors and represents a universal liability for this class of compounds as they advance into the clinic.

## Discussion

Using AML as a model system for response and resistance, we found that acute CBP/p300 HAT inhibition leads to genome-wide histone hypoacetylation and consequent loss of enhancer-mediated leukemic transcription. Interestingly, these transcriptional effects are substantively identical in hypersensitive and intrinsically resistant AML cell lines, ultimately indicating that this primary, mechanistic response is not predictive of downstream anti-proliferative effects. Instead, secondary adaptations to the initial transcriptional response vary dramatically among AML cell lines and are likely more important as a reflection, and possibly even a determinant, of sensitivity. Our discovery that the primary transcriptional responses to drug treatment are not predictive of an anti-proliferative effect has also been observed for BET bromodomain inhibition^32^, suggesting that it may be a general feature of cellular responses to drugs targeting transcriptional regulators.

By modeling acquired resistance to HAT inhibition, our study revealed an unexpected ability of cells to interrupt drug-target engagement by elevating the production of a competitive-binding, endogenous co-factor. This is an unprecedented mechanism of resistance that has not been described for other metabolite-competitive compounds (e.g. ATP-competitive kinase inhibitors). For other emerging drug classes, such as SAM-competitive methyltransferase inhibitors, it is possible that analogous resistance mechanisms are relevant and should be a new source of vigilance. The facile acquisition of genetic alterations that can alter CoA metabolism to overcome the negative selective pressure of HAT inhibition reflects the surprisingly malleable nature of this biosynthetic pathway. Our work therefore points to a major new consideration for the optimization of this class of drugs, which is distinct from the widely appreciated link between cellular metabolism and histone acetyltransferase activity^46–51^. In our panel of AML cell lines, the concentration of acetyl-CoA in drug-naïve populations did not likely influence intrinsic responses, as the dose-responsive inhibition of CBP/p300 did not differ between intrinsically sensitive and insensitive AML cell lines. However, we cannot rule out the possibility that acetyl-CoA production may determine responses in other indications or even influence on-target toxicity in normal tissues.

We have demonstrated that changes in acetyl-CoA concentrations can arise through alterations to *PANK3, PANK4*, and *SLC5A6*. However, given the apparent tolerance of cells to elevated CoA production, it seems likely that additional mechanisms to modulate this pathway await discovery. Our data suggest that *PANK3*, in particular, is highly permissive to hypermorphic mutations, since the simple obstruction of feedback inhibitory acetyl-CoA binding is sufficient to drive runaway CoA production, even if such mutations also impair intrinsic catalytic activity. The cellular CoA/acetyl-CoA concentration is determined by the expression level and feedback inhibition of all three PANK isoforms^6,38^. Thus, elevated cellular CoA may also arise from hypermorphic mutations within the identical active sites of other PANK isoforms or through their transcriptional upregulation. We might also expect sensitivity to HAT inhibitors would be modulated by cell extrinsic factors that are known to regulate intracellular acetyl-CoA concentrations, such as the availability of glucose^52^.

We foresee several strategies to overcome HAT inhibitor resistance, including (1) allosteric inhibitors, which would not be competitive in binding with acetyl-CoA, (2) targeted protein degradation, which may be able to overcome competition through its catalytic and irreversible effects, (3) covalent inhibition, and (4) leveraging synthetic vulnerabilities that may accompany the dysregulation of CoA metabolism. Allosteric and covalent inhibitors have been reported for CBP/p300^37,53^, but they remain unexplored for other HAT targets. Furthermore, bromodomain inhibitors only partially disrupt CBP/p300 catalytic activity^54^ and may therefore not present a viable strategy for all indications in which elevated acetyl-CoA production mediates resistance to HAT inhibition. Finally, since this resistance applies to all acetyl-CoA-competitive HAT inhibitors, a target-independent mechanism to overcome or circumvent resistance would be of substantial value. We expect that the diversion of cellular resources to elevate the production of CoA, as well as the downstream metabolic consequences of this change, may create new vulnerabilities in the cell that could be exploited for sequential or concurrent combination therapies. These and other strategies will prove essential for the design and optimization of next-generation therapeutics targeting acetyltransferases.

## Acknowledgements

We gratefully acknowledge B.F. Cravatt for critical feedback of the manuscript while it was in preparation. We thank Karen Miller for protein biochemistry, Katie Creed for cell culture and Matthew Frank for mass spectrometry. Sequencing was performed by the Scripps Research Next Generation Sequencing Core (La Jolla). This work was supported by the National Institutes of Health (NIH) through an NIH Director’s Early Independence Award (DP5-OD26380) and by the Ono Pharma Foundation (M.A.E.). This work was also supported by Cancer Center Support Grant CA21765 and the American Lebanese Syrian Associated Charities (C.O.R.).

## Methods

### Cell lines and lentivirus production

MOLM-13 (RPMI 1640 supplemented with 10% Fetal Bovine Serum (FBS) and 1x Antibiotic-Antimycotic (anti-anti, Gibco), MV4;11 (RPMI 1640 supplemented with 10% FBS and 1x anti-anti), OCI/AML-2 (RPMI 1640 supplemented with 10% FBS and 1x anti-anti), HL-60 (RPMI 1640 supplemented with 20% FBS and 1x anti-anti), NOMO-1 (RPMI 1640 supplemented with 20% FBS and 1x anti-anti), and SKM-1 (RPMI 1640 supplemented with 20% FBS and 1x anti-anti) were provided by the laboratory of Prof. James E. Bradner. RPMI-8226 (RPMI supplemented with 15% FBS and 1x anti-anti) cells were purchased from DSMZ. Production of lentivirus was performed in Lenti-X 293T cells (Takara) by co-transection with pMD2.G, psPAX2 (Addgene #12259 and 12260, respectively), and a given lentiviral expression plasmid using polyethylenimine. Supernatants containing lentivirus were collected at 48 and 72 hours after transfection, filtered through a 0.22 μM membrane, and concentrated 20-fold with Lenti-X Concentrator (Takara). Cell lines were transduced by spinoculation at 500 g for 1 hour at room temperature with 8 μg/mL polybrene (EMD Millipore). HEK293T cells (ATCC, #CRL-3216) were maintained in Dulbecco’s modified Eagle’s medium (DMEM, Bio Whittaker/Lonza) supplemented with 10% FBS, 2 mM L-glutamine and 50 U/ml penicillin/streptomycin. Cell lines generated in this study are available from authors upon request.

### Immunoblotting

For histone blots, equal numbers of treated cells were collected by centrifugation and washed twice with ice-cold PBS. Cell pellets were then lysed in PBS containing 0.5% Triton X (v/v) and 1x Halt Protease Inhibitor Cocktail (Thermo) on ice for 15 minutes. Lysates were centrifuged at 2000 rpm for 10 minutes at 4° C and supernatant was discarded. Pellets were washed again with lysis buffer followed by centrifugation and removal of supernatant. Pellets were then resuspended in 0.2 N HCl at a density of 4×10^7^ cell per mL and acid extracted overnight at 4° C. Samples were then centrifuged, and supernatant was collected. Whole cell lysates were prepared by solubilizing cell pellets in CelLytic M (Sigma Cat# C2978), 1X Halt Protease Inhibitor cocktail (Thermo) and 0.1% Benzonase (EMD Millipore) and lysing on ice for 15 minutes. Cleared lysates were prepared by centrifugation at 18,000xg for 10 minutes at 4° C. 1x SDS Sample Buffer and 2.5% 2-mercaptoethanol were added to all samples before separation on Bis-Tris gels. Loading of equal amounts of protein was ensured by a BCA protein assay kit (Pierce). Protein was transferred to nitrocellulose membrane and the membrane was blocked for 1 hour in 5% non-fat milk in 1x TBS-T before incubation with primary antibodies overnight at 4 °C. Primary antibodies were detected with infrared secondary antibodies (IRDye) on the Odyssey CLx Images (LI-COR). Quantitation of blots was performed using ImageStudio Lite (LI-COR).

### Cellular viability and proliferation assays

For viability assays, 1,000 cells were plated into 384-well plates (Greiner, Cat# 07-000-056) in 50 μL of media. 100 nL of compounds in DMSO were pinned into the plates using a Biomek FX Automated Workstation (Beckman Coulter). After 72 hours, 25 μL of ATPLite reagent was added to each well. Luminescence was detected with an Envision multilabel plate reader (PerkinElmer, Model No. 2104). For cell growth assays, cells were seeded at 20,000 cells/well in a 96-well plate with compound added as 1:1000 dilutions of DMSO stocks. Cell number was determined every 3-4 days using the Countess automated cell counter (Invitrogen). 20,000 cells were then re-plated with fresh media and compound. Each treatment was done in triplicate. Cumulative cell count was determined by back calculation.

### Cell cycle analysis

Cells were seeded at 200,000 cells/well in a 48-well plate with compound added as 1:1000 dilutions of DMSO stocks. Cell cycle analysis was performed with FITC BrdU Flow Kit (BD Pharmingen, Cat# 559619) according to manufacturer’s protocol. Briefly, treated cells were labeled for 30 minutes with 10 μM BrdU. Cells were then permeabilized and fixed using kit buffers. To expose BrdU epitopes, cells were incubated with 300 μg/mL DNase at 37C for 1 hour. Cells were then washed and incubated with a FITC conjugated anti-BrdU antibody for 20 minutes at room temperature. Cells were then washed and incubated in 7-AAD solution. Finally, cells were resuspended in FACS staining buffer (3% FBS in PBS) and analyzed by flow cytometry on a Novocyte 3000 (Agilent). Flow cytometry results were analyzed using NovoExpress (version 1.5.6).

### Chromatin immunoprecipitation followed by DNA sequencing with spike-in normalization (ChIP-Rx)

ChIP-Rx was performed as previously described. For H3K18ac and H3K27ac ChIPs 50 million cells were treated per condition. For ENL and BRD4 ChIPs 100 million cells were treated. For RNA Polymerase II ChIPs 150 million cells were treated per condition. For histone PTM ChIPs crosslinked Drosophila S3 cells were used for spike-in normalization and for all other ChIPs crosslinked mouse NIH/3T3 cells were used for spike-in normalization. Treated cells were crosslinked by adding a 1/10^th^ volume of 10x crosslinking solution (11% formaldehyde, 50 mM HEPES pH 7.3, 100 mM NaCl, 1 mM EDTA pH 8.0, 0.5 mM EGTA pH 8.0). After 10 minutes, crosslinking was quenched with 125 mM Glycine. Crosslinked cells were washed three times in ice-cold 1x DPBS pH 7.4 and then flash frozen in liquid nitrogen and stored at −80° C. Pellets were resuspended in cold lysis buffer 1 (50 mM HEPES pH 7.3, 140 mM NaCl, 1 mM EDTA, 10% glycerol, 0.5% NP-40, and 0.25% Triton X-100, 1X HALT protease inhibitor cocktail) and rotated for 10 minutes at 4C. Crosslinked spike-in cell pellets were resuspended in lysis buffer 1 and equal volumes were added to samples at a ratio of 1 spike-in cell to 10 sample cells for mouse spike-ins and 1 spike-in cell to 5 sample cells for drosophila spike-ins. Pellets were collected by centrifugation at 1350 g before being resuspended in cold lysis buffer 2 (10 mM Tris-HCl pH 8.0, 200 mM NaCl, 1 mM EDTA pH 8.0 and 0.5 mM EGTA pH 8.0, 1X HALT protease inhibitor cocktail) and rotated again for 10 minutes at 4° C. Pellets were again collected by centrifugation at 1350 g before being resuspended at ~30 million cells/mL in sonication buffer (50 mM HEPES pH 7.3, 140 mM NaCl, 1 mM EDTA, 1 mM EGTA, 1% Triton X-100, 0.1% Na-deoxycholate, 0.5% SDS, 1X HALT protease inhibitor cocktail) and split into 250 μL volumes in 1.5 mL Bioruptor Plus TPX microtubes (Diagenode, #C30010010). Cells were sonicated at 4° C for 20 minutes using a waterbath sonicator cycling 30 seconds on and 30 seconds off (Bioruptor Pico, Diagenode). Sonicated lysates were cleared by centrifugation at 18,000 g for 10 minutes at 4° C. Lysates from the same sample were pooled and 50 μL was set aside and stored at −20° C as the input. The remaining lysates were diluted 1:5 in sonication buffer without SDS. Antibody-bound magnetic protein G beads (Dynabeads, Thermofisher Scientific) were added to the sonicated chromatin supernatant and rotated overnight at 4° C. Used 100 μL of beads with 10 μL anti-H3K27ac (Abcam, ab4729), 20 μL anti-H3K18ac (EMD Millipore, #07-354), 20 μL anti-RNAPII (Diagenode, #C15100055), 10 μg anti-ENL (Cell Signaling Technology, #14893S), and 10 μg anti-BRD4 (Bethyl Laboratories, # A301-985A100). Beads were then washed twice with 1 mL cold sonication buffer, once with 1 mL cold sonication buffer supplemented with 500mM NaCl, once with cold LiCl wash buffer (20 mM Tris pH 8.0, 1 mM EDTA, 250 mM LiCl, 0.5% NP-40, 0.5% Na-deoxycholate), and once with TE supplemented with 50 mM NaCl. Beads were then resuspended in 215 μL of elution buffer (50 mM Tris-HCl pH 8, 10 mM EDTA, and 1% SDS). Chromatin was eluted by vortexing the beads every 5 minutes while heating at 65° C for 15 minutes. Beads were centrifuged at 20,000 g for 30 seconds and 200 μL of supernatant was collected and moved to a new tube. Eluted chromatin and input samples were incubated overnight at 65° C to reverse crosslinks. RNA was digested with 0.2 mg/mL RNAse A at 37° C for 2 hours. Protein was digested with 0.2 mg/mL Proteinase K at 55°C for 30 minutes. DNA was isolated with MinElute PCR Purification Kit (Qiagen). Libraries for Illumina sequencing were prepared using the ThruPLEX DNA-seq Kit (Takara) and DNA Single Index Kit - 12S Set A (Takara) according to manufacturer’s protocol. Samples were then multiplexed and sequenced on an Illumina NextSeq 500 (singleend 75 bp reads).

Sequencing reads were aligned to human genome build Hg19 and spike-in reference genomes (Mm9 or Dm3) by Bowtie2 (v2.2.9) with all default settings except for -N 1 (allowing 1 mismatch in seed alignment). SAM alignment files were compressed to BAM files, sorted and indexed with Samtools (v1.9). Peaks were called using the model-based analysis of ChIP-seq (MACS, v1.4.2) with ChIP input samples as background and a P value threshold of 1e-9. Annotation of putative enhancers and super-enhancers and their predicted target genes was performed using Rank Ordering of Super Enhancers (ROSE2) analysis of H3K27ac ChIP-seq data with default parameters to create a GFF file. A GFF file for promoters was created by filtering for all annotated transcription start sites (TSS) in Hg19 that overlapped with an H3K27ac peak and defining the promoter as +/- 1 kb from the TSS. H3K27ac and H3K18ac read density (reads/bp) at these sites was calculated using Bamliquidator (v1.0). For BRD4 and ENL, the GFF files were further filtered to only include sites that overlapped with a BRD4 or ENL peak before read density was calculated. Read density values were then normalized by dividing by the total number of reads that aligned to the spike-in reference genome producing a reference-adjusted reads per million (RRPM). For RNA Pol II ChIP analysis, GFFs at all genes in Hg19 were made for the TSS (+/- 300 bp) and the gene body (TSS + 300 bp to the transcription termination region + 3 kb) and these were filtered to include only genes where the TSS overlapped with an RNA Pol II peak. Read density at these regions was calculated and normalized as described above. The traveling ratio of RNA Pol II was calculated by dividing the TSS signal by the gene body signal.

### 3’ mRNA-seq with ERCC RNA Spike-In Mix normalization

For 3’ mRNA-seq analysis of gene expression, cells were treated in triplicate with A-485 (3 μM) or DMSO (0.1%) for 48 hours. After treatment, cells were counted using the Countess automated cell counter and RNA was isolated with the Qiagen RNeasy miniprep kit. ERCC RNA Spike-In Control SIRV Set 3 was spiked into cell number normalized volumes of eluted RNA. Libraries for Illumina sequencing were prepared using the QuantSeq 3’ mRNA-Seq Library Prep Kit FWD for Illumina (Lexogen, Cat# 015) and multiplexed libraries were sequenced on Illumina NextSeq 500 (single end 75 bp reads). SlamDunk (with the -q option to ignore SLAM-seq scoring) was used to align reads to human genome build hg19 and ERCC SIRV Set 3 and generate count tables with counts per million (CPM) at all 3’ UTRs using SlamDunk. Locally estimated scatterplot smoothing (LOESS) normalization was performed for spike-in normalization by using the normalize.loess function of the affy v1.50.0 R package76. The LOESS distribution was fit on the SIRV-Set-3 spike-ins and then applied to the whole dataset. Average log2-transformed CPM fold change values were calculated from the means of triplicate treatments. For each cell line, any transcripts with a maximum average CPM < 3 in the DMSO and A-485 treated groups were excluded from analysis in that cell line. Gene set enrichment analysis was performed using GSEA software (Broad Institute) against all gene sets in MSigDB collection C2.

### Thiol(SH)-linked alkylation for the metabolic sequencing of RNA (SLAM-seq)

For SLAM-seq analysis of gene expression, cells were pre-treated in triplicate with 3 μM A-485 or 0.1% DMSO for 1 hour. 100 μM 4-thiouracil was then added to label newly synthesized RNAs for 1 hour. While limiting exposure to light, cells were collected by centrifugation and RNA was isolated with the RNeasy miniprep kit (Qiagen). RNA yield was quantified fluorometrically with the Qubit RNA XR Assay kit and 5 μg of RNA from each sample was taken for alkylation and brought to a volume of 15 μL. RNA was alkylated by addition of 5 μL 100 mM iodoacetamide, 25 μL DMSO, and 5 μL of sodium phosphate buffer (0.5 M, pH 8) and incubating at 50C for 15 minutes. Reactions were then quenched by addition of 1 μL 1M DTT. RNA was purified by ethanol precipitation and again quantified fluorometrically. Libraries for Illumina sequencing were prepared using the QuantSeq 3’ mRNA-Seq Library Prep Kit FWD for Illumina (Lexogen, Cat# 015) and multiplexed libraries were sequenced on Illumina NextSeq 500 (single end 75 bp reads). SlamDunk with default parameters was used to align reads to human genome build hg19, calculate CPM and T>C conversion rate at all 3’ UTRs. Average log2-transformed CPM and conversion rate fold change values were calculated from the means of triplicate treatments. For each cell line, any transcripts with a maximum average CPM < 3 among treatment groups were excluded from analysis in that cell line. Grouping of genes into those predicted to be regulated by no enhancer, typical enhancers, and super-enhancers was done using predictions from ROSE2 as described above.

### CRISPR-Cas9 competition assays

Cell lines were engineered to express Cas9 by transduction with a lenti-Cas9-Blast (Addgene #52962) lentivirus and selection with 10 μg/mL blasticidin. Cas9-expressing cells were transduced in triplicate with a Lenti_sgRNA_EFS_GFP (Addgene #65656) lentivirus that co-expressed an sgRNA and GFP in 96-well plates. Transduction efficiency was determined by flow cytometry 3 d posttransduction and the experiment was continued if the transduction efficiency was between 30-60%. For sgRNA dropout experiments with CREBBP- and EP300-targeting sgRNAs the proportion of GFP+ cells was monitored every 3-4 days by flow cytometry. For competitive growth assays with drug selection, the proportion of GFP+ cells was monitored every 7 days by flow cytometry. sgRNA sequences used for these experiments are as follows: Luciferase 5’-CCCGGCGCCATTCTATCCGC-3’; RPS19 5’-GTAGAACCAGTTCTCATCGT-3’; EP300-sg1 5’-ATGGTGAACCATAAGGATTG-3’; EP300-sg2 5’-CTGTAATAAGTGGCATCACG-3’; EP300-sg3 5’-GGTACGACTAGGTACAGGCG-3’; EP300-sg4 5’-GTGGCACGAAGATATTACTC-3’; CREBBP-sg1 5’-ATTGCCCCCCTCCAAACACG-3’; CREBBP-sg2 5’-CAGGACGGTACTTACGTCTG-3’; CREBBP-sg3 5’-CCGCAAATGACTGGTCACGC-3’; CREBBP-sg4 5’-CTTAGCCCACTGATGAACGA-3’; PANK4-sg1 5’-GGAATCCTTCTGGTACACGA-3’; PANK4-sg2 5’-TTTGGAGGCTTCTTTATCCG-3’; SLC5A6-sg1 5’-CATGATGCTCTCCTTATACG-3’; SLC5A6-sg2 5’-CCTGCATTGAGAGCCAATGA-3’.

### Generation of knockout and CRISPR base editor populations

To create knockout populations, Cas9-expressing cells were transduced with a Lenti_sgRNA_EFS_GFP (Addgene #65656) lentivirus co-expressing an sgRNA and GFP to achieve an efficiency >95% as confirmed by flow cytometry. For PANK4 and SLC5A6 knockouts clonal populations were generated from the polyclonal pool by limiting dilution. Clonal knockouts were confirmed by immunoblot for the corresponding protein or through confirmation of genetic disruption by TIDE. For base-editing, cells were transduced with a pRDA_256 (Addgene #158581) co-expressing an sgRNA and BE3.9max with a puromycin resistance cassette. Transduced cells were selected for with puromycin (2 μg/mL) and clonal populations were generated by limiting dilution. Presence of the intended base edit was confirmed by sanger sequencing. For knockouts of PANK4 and SLC5A6 the same sgRNAs used in the competitive growth assay were used. For knockout of PANK3 the sgRNA was as follows: PANK3-sg 5’-ACAACTATAAACGAGTGACT-3’. For base editor experiments the guides used were as follows: AAVS1 5’-GGGGCCACTAGGGACAGGAT-3’; EP300-BE-sg1 5’-GTCCACCATGTGCATGCTGG-3’; EP300-BE-sg2 5’-GTGGTCCACCATGTGCATGC-3’.

### Genome-wide sgRNA screening

250 million Cas9-expressing MOLM-13 were transduced with the Brunello sgRNA library (Addgene #73179) at a MOI around 0.3, aiming for 1000-fold representation of each guide. Transduced cells were selected with 2 μg/mL puromycin and allowed to triple before collecting an initial 80 million cell pellet to measure initial sgRNA representation and splitting the remaining cells into treatment arms: 250 nM A-485 (later increased to 300 nM) and vehicle. Treated populations were cultured for 21 days and 80 million cell pellets were collected for each treatment arm. Genomic DNA was extracted using Quick-DNA Midiprep Plus Kit (Zymo Research, #D4075). Illumina sequencing libraries were prepared by first amplifying the sgRNA containing vector sequences from the gDNA and then using a second PCR reaction to append sequencing adapters^55^. Libraries were sequenced on Illumina NextSeq 500 (single end 75 bp reads). Sequencing reads were trimmed using cutadapt (v1.7) to remove vector derived sequences and all analysis was performed using MaGeCK^56^. Briefly, the MaGeCK count function was used to align reads to the Brunello sgRNA library and generate read count tables for all samples. The MaGeCK MLE function was then used with default parameters to compare sgRNA abundance changes in DMSO and A-485 samples and create a gene-level beta score indicative of the relative enrichment or depletion of sgRNAs targeting that gene. Screening data from MM1S cells was obtained from a previous study^18^ (GEO ID: GSM5278023-GSM5278026) and the MaGeCK count function was used to align reads to the Brunello and GeCKOv2 sgRNA libraries. To generate similar plots to the original report, the MaGeCK RRA function was used to determine gene-level sgRNA enrichment and the *P* values for this enrichment in two distinct replicates were plotted against each other.

### Generation of evasive resistant cell line

To generate a resistant cell line, MOLM-13 cells were serially replated in media containing either vehicle (0.1% DMSO) or increasing concentations of A-485. Starting from 50 nM A-485, the dose was increased with each passage by about 10% until a maximum dose of 2 μM was reached (MOLM-13-2R). These cells were cultured in media containing 2 μM A-485 until it was verified that they maintained resistance to A-485 without constant exposure, at which point they were cultured in media without any drug.

### Whole-exome sequencing

Genomic DNA was isolated from cell lines using DNeasy Blood & Tissue Kit (Qiagen). Whole-exome sequencing was performed using SureSelect Human All Exon V7 (Agilent) according to manufacturer’s protocol and sequenced on an Illumina NextSeq 500 (paired-end 150 bp reads). Reads were aligned to human genome build hg19 with bwa (version 0.7.12). Aligned reads were sorted and indexed with samtools (version 1.10). Somatic variants in the evasive resistant cell lines were detected and filtered using GATK Mutect2 and FilterMutectCalls (GATK version 4.1.9.0) using the wild-type MOLM-13 cell line exome as a matched normal. GATK Funcotator was used to annotate detected variants. Annotated variant calls that passed GATK filtering were further filtered to include only variants that met the following criteria: read depth >= 50, variant allele fraction >= 0.1, and variant allele predicted to cause a coding mutation.

### Ectopic expression of PANK3

PANK3 mutations were made in a gateway compatible pDONR plasmid containing a PANK3 ORF (a gift from the lab of Ben Cravatt) by amplifying the whole plasmid with primers containing the desired mutations and using HiFi DNA Assembly Master Mix (NEB, # E2621) to re-circularize the amplicon. LR Clonase II (ThermoFisher, # 12538120) reactions were used to move the PANK3 ORFs into the lentiviral expression vector pLEX_307 (Addgene #41392). Cells were transduced with lentivirus made from these vectors, selected for with puromycin (2 μg/mL), and expression of PANK3 was confirmed by immunoblot for the V5 tag. For experiments in HEK293T, cells were transfected using FuGENE® 6 reagent (Promega) combined with 12 μg of the indicated plasmid cDNA construct: pcDNA3.1(-) vector or vector expressing either PANK3 or PANK3_G193A_. Cells were grown for 48 hours and harvested for extraction of total CoA and acetyl CoA.

### Protein thermal shifts

Protein thermal shift experiments were conducted in a mixture containing 100mM HEPES, pH 7.0, 25x Sypro Orange dye, 10 mM MgCl_2_, 5 μM PANK3 or PANK3(G193A), and acetyl CoA or ATP at the indicated concentrations. Reaction mixture (30 μl) was aliquoted in triplicate into optically clear 96-well PCR plates, centrifuged and placed in an ABI 7500 Fast real time PCR system. The temperature was ramped from 25 °C to 95 °C at a rate of 1 °C/min, and fluorescence was measured using a TAMRA filter set (Ex 560 nm and Em 582 nm). The fluorescence intensity as a function of temperature was fit to the first derivative of the Boltzman sigmoidal equation using GraphPad software to determine the temperature corresponding to the denaturation of 50% of the protein. Each experiment was performed in triplicate and the melting temperatures were the average of each set.

### PANK activity assays

PANK assays were performed as described (PMID: 30352999, PMID: 34524863, PMID: 27555321). Briefly, PANK3 activity assays were performed in a reaction mixture that contained 100 mM Tris-HCl, pH 7.5, 10 mM MgCl_2_, 2.5 mM ATP, 45 μM D-[1–^14^C]pantothenate (specific activity, 22.5 mCi/mmol) and 150 ng of PANK3 or 8 μg of PANK3(G193A). PANK3 concentrations were calculated using the extinction coefficient at 280 nm of 39,225 M-1 cm-1. The assay was linear with time and protein. After 10 min at 37 °C, the reaction was stopped by the addition of 4 μl of 10% (v/v) acetic acid. The mixture was spotted onto a DE81 disk, washed with three successive changes of 1% acetic acid in 95% ethanol and product formation was determined by scintillation counting of the dried disc. The reaction mixes for the determination of the acetyl-CoA IC_50_ contained 100 mM Tris-HCl, pH 7.5, 10 mM MgCl_2_, 1 mM ATP, 45 μM D-[1-^14^C]pantothenate (specific activity 22.5 mCi/mmol) and 150 ng of PANK3 or 8 μg of PANK3(G193A). All the experiments were performed twice in duplicate, and the data are reported as an average ± data range.

### Total CoA determination

Total CoA was determined as described earlier (PMID: 30352999, PMID: 31609347). Approximately 6 X 10^6^ cultured cells were washed with ice-cold phosphate-buffered saline that was quickly aspirated off the dish, and cells were washed again with ice-cold water that was aspirated off to remove residual saline. Ice-cold water (1 ml) was added to the culture dish, cells were scraped into the cold water and the cell suspension was transferred to a glass test tube containing 400 μl of 0.25 M KOH and 1.5 ml water. Samples were incubated for 2 hours at 55 °C to hydrolyze CoA thioesters, the pH was lowered to 8 by addition of 1 M Tris base and monobromobimane (mBBr, Life technologies) was added in excess to derivatize the CoASH. The CoA-bimane product was purified on a SPE 2-(2-pyridyl)ethyl column and fractionated by HPLC with fluorescent detection λex = 393 nm, λem = 470 nm. The retention time was determined by running a CoA-bimane standard before each set of samples. Quantification was done using a calibration curve made with the CoA-bimane standard.

### Acetyl-CoA measurement by mass spectrometry

Acyl-CoAs were quantified as described earlier (PMID: 31609347, PMID: 34524863). Briefly, approximately 6 × 10^6^ MOLM-13 cells were resuspended in 2 ml methanol and 1 ml water and incubated on ice for 30 minutes. Chloroform 1.5 ml and 1.2 ml water were added to remove lipids. The aqueous top layer was collected and 30 pmol of [^13^C_2_]acetyl-CoA internal standard (Sigma) was added and dried overnight in a Savant Speedvac Concentrator SPD 1010 (Thermo-Fisher). Samples were injected onto an Acquity UPLC HSS C18, 2.5 μm, 3.0 × 150 mm column at 40°C (Waters) using a flow rate of 0.2 ml/min. Solvent A was 10 mM ammonium acetate, pH 6.8, and Solvent B was 95% acetonitrile + 10 mM ammonium acetate, pH 6.8. The HPLC program was: starting solvent mixture of 95% A/5% B, 0 to 2 min isocratic with 5% B; 2 to 20 min linear gradient 100% B; 20 to 25 min isocratic with 100% B; 25 to 27 min linear gradient to 5% B; 27 to 31 min isocratic with 5% B. The Sciex QTrap 4500 was operated in the positive mode, and the ion source parameters were: ion spray voltage, 5500 V; curtain gas, 15 psi; temperature, 400 °C; collision gas, medium; ion source gas 1, 15 psi; ion source gas 2, 20 psi; declustering potential, 60 V; and collision energy, 45 V. The MRM transitions for acetyl-CoA, 810.1/303 was used. was used to calculate relative abundance of the acyl-CoAs in cells. The system was controlled by the Analyst software (Sciex) and analyzed with MultiQuant 3.0.2 software (Sciex). Peaks corresponding to acetyl-CoA species were quantified relative to the [^13^C_2_]acetyl-CoA internal standard.

### Data and Code Availability

The sequencing data in this publication have been deposited in the NCBI Gene Expression Omnibus (GEO) under accession number GSE211054. The software and code used to analyze this data is all available online and has been detailed in the Methods section.

**Extended Data Figure 1:**
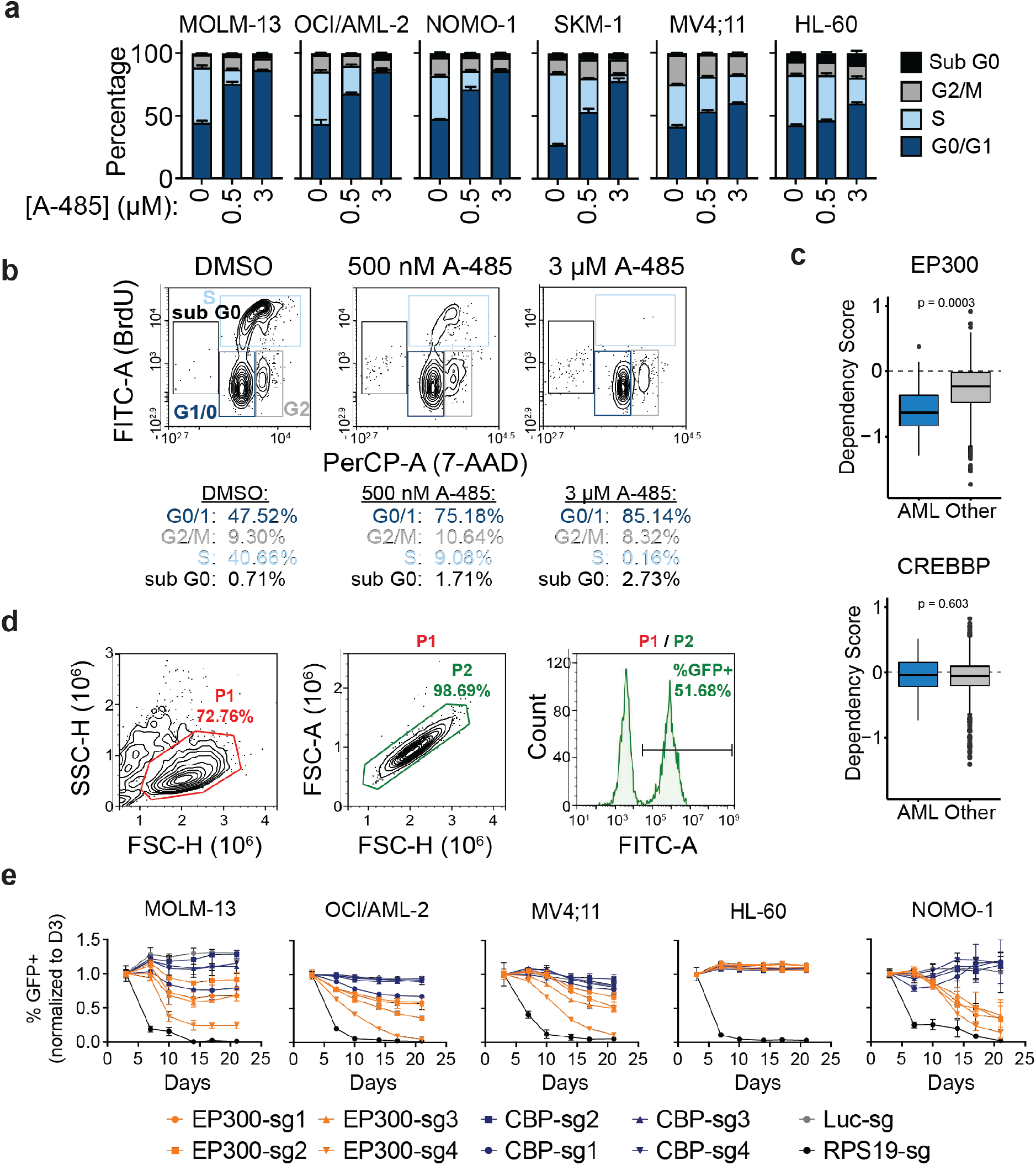
Response of AML cell lines to CBP/p300 disruption is heterogenous. **a**, Cell cycle analysis in panel of AML cell lines following 48-h treatments by BrdU incorporation. Mean ± SD. **b**, Representative flow cytometry plots for cell cycle analysis in MOLM-13 following 48-h treatments. **c**, Boxplots of dependency scores of EP300 and CREBBP in AML and all other cancer lineages. Data from Cancer Dependency Map release 22Q1. P values were determined by two-tailed Student’s t-test. **b**, Representative flow cytometry plots of gating strategy used for CRISPR-Cas9-based competitive growth assay. **e**, CRISPR-Cas9-based competitive growth assay in panel of AML cell lines with 4 EP300-targeting sgRNA (light blue), 4 CREBBP-targeting sgRNA (dark blue), a luciferase-targeting sgRNA negative control (grey), and a RPS19-targeting sgRNA positive control (black). Mean ± SD, n = 3.

**Extended Data Figure 2:**
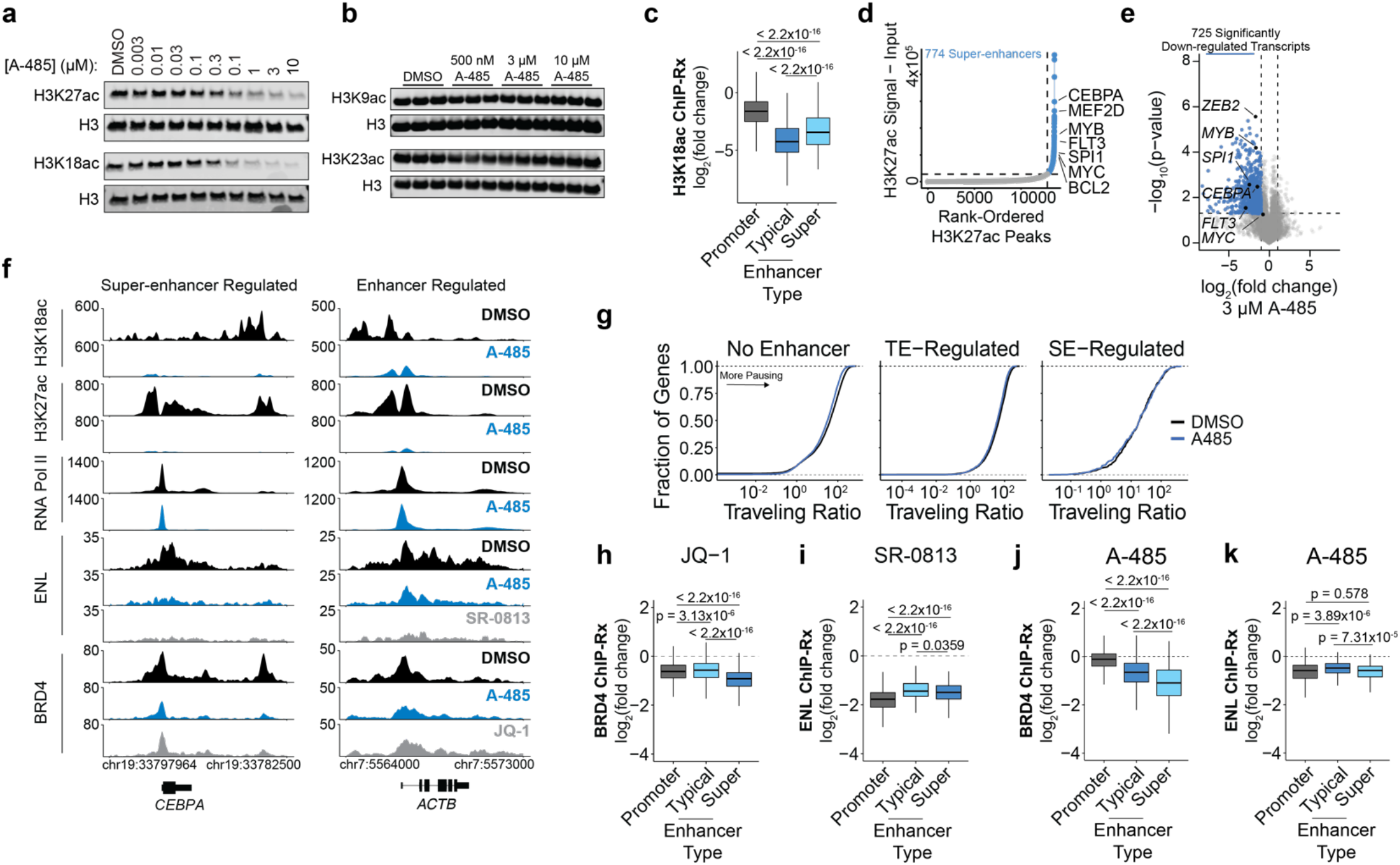
Acute transcriptional response to CBP/p300 HAT inhibition in MOLM-13. **a**, Immunoblot of MOLM-13 treated with A-485 for 2 h and probed for total H3 and CBP/p300 substrates H3K27ac and H3K18ac. **b**, Immunoblot of MOLM-13 treated with A-485 and probed for total H3 and non-CBP/p300 substrates H3K9ac and H3K23ac. **c**, Boxplots of change in H3K18ac ChIP-Rx signal at promoters (n = 12650), typical enhancers (n = 12823) and superenhancers (n = 774). **d**, Rank-ordered plot of H3K27ac ChIP-Rx signal in MOLM-13 at promoter-distal H3K27ac peaks used to define super-enhancers. **e**, Volcano plot of DMSO-normalized changes in nascent transcription as measured by SLAM-seq following 2-h of treatment with A-485 (3 μM) in MOLM-13. *P* value calculated with two-tailed Student’s t-test (n = 3). **f**, Gene tracks of ChIP-Rx signal (RRPM) in MOLM-13 at representative loci. **g**, Cumulative distribution plot of RNA Pol II traveling ratio at genes predicted to be regulated by no enhancer (n = 11028), typical enhancers (n = 3960), and super-enhancers (n = 451) determined by RNA Pol II ChIP-Rx after 2-h of A-485 (3 μM) treatment. **h**, Boxplots of DMSO-normalized changes in BRD4 abundance at BRD4-bound promoters (n = 10720), typical enhancers (n = 6262), and superenhancers (n = 774) following 2-h of JQ-1 treatment (300 nM). **i**, Boxplots of DMSO-normalized changes in ENL abundance at ENL-bound promoters (n = 856), typical enhancers (n = 357), and super-enhancers (n = 179) following 2-h of SR-0813 (10 μM) treatment. **i**, As in **h**, but following 2-h of A-485 treatment (3 μM). **k**, As in **i**, but following 2-h of A-485 treatment (3 μM). For all boxplots, boxes represent 25-75 percentiles with whiskers extending to 1.5 * the interquartile range (IQR) and P values were determined by a two-tailed Welch’s t-test.

**Extended Data Figure 3:**
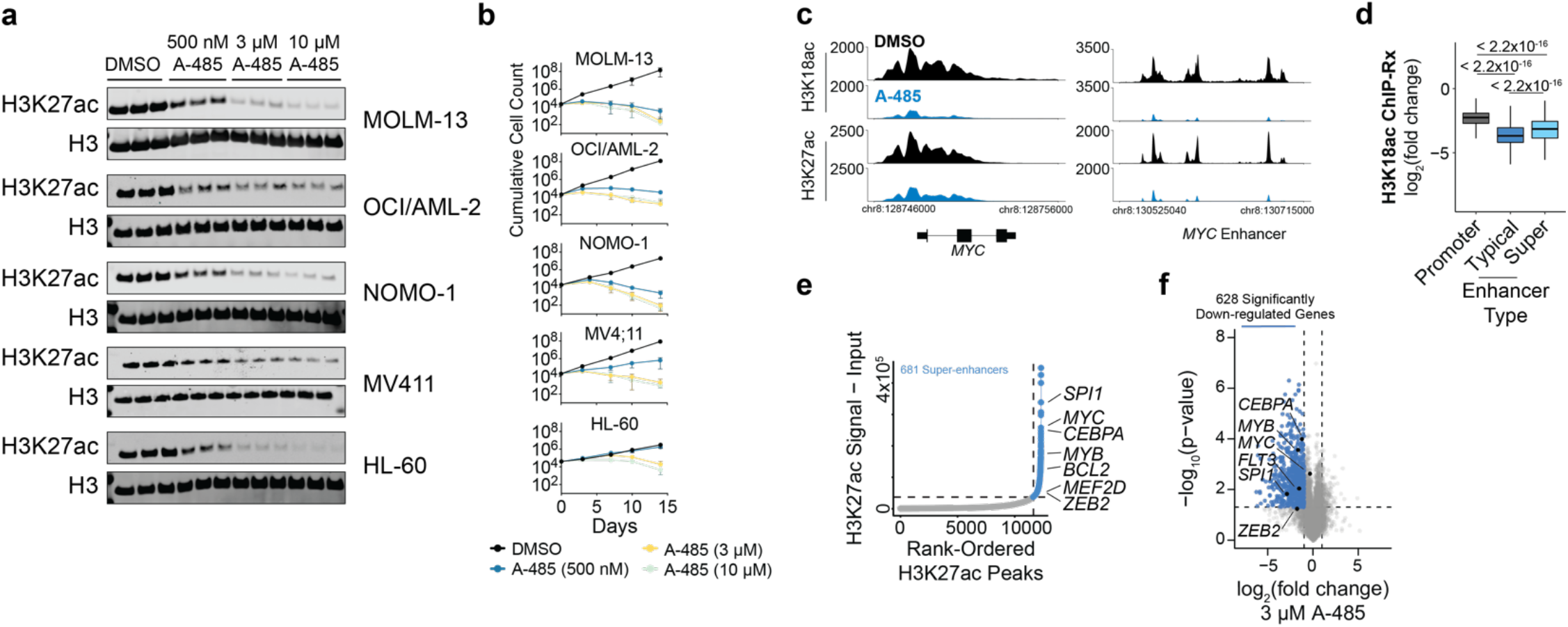
Acute transcriptional responses to CBP/p300 HAT inhibition in HL-60 and other AML cell lines. **a**, Immunoblot for H3K27ac and total H3 in panel of AML cell lines following 2-h of A-485 treatment. **b,** Proliferation of AML cell lines in response to A-485 treatment over 14 days. Mean ± SD, n = 3. **c**, Gene tracks of ChIP-Rx signal (RRPM) in HL-60 at representative loci. **d,** Boxplots of change in H3K18ac ChIP-Rx signal in HL-60 at promoters (n = 12171), typical enhancers (n = 11635) and super-enhancers (n = 681). **g**, Rank-ordered plot of H3K27ac ChIP-Rx signal in HL-60 at promoter-distal H3K27ac peaks used to define super-enhancers. **f,** Volcano plot of DMSO-normalized changes in nascent transcription as measured by SLAM-seq following 2 hours of treatment with A-485 (3 μM) in HL-60. *P* value calculated with two-tailed Student’s t-test (n = 3). For all boxplots, boxes represent 25-75 percentiles with whiskers extending to 1.5 * the interquartile range (IQR) and P values were determined by a two-tailed Welch’s t-test.

**Extended Data Figure 4:**
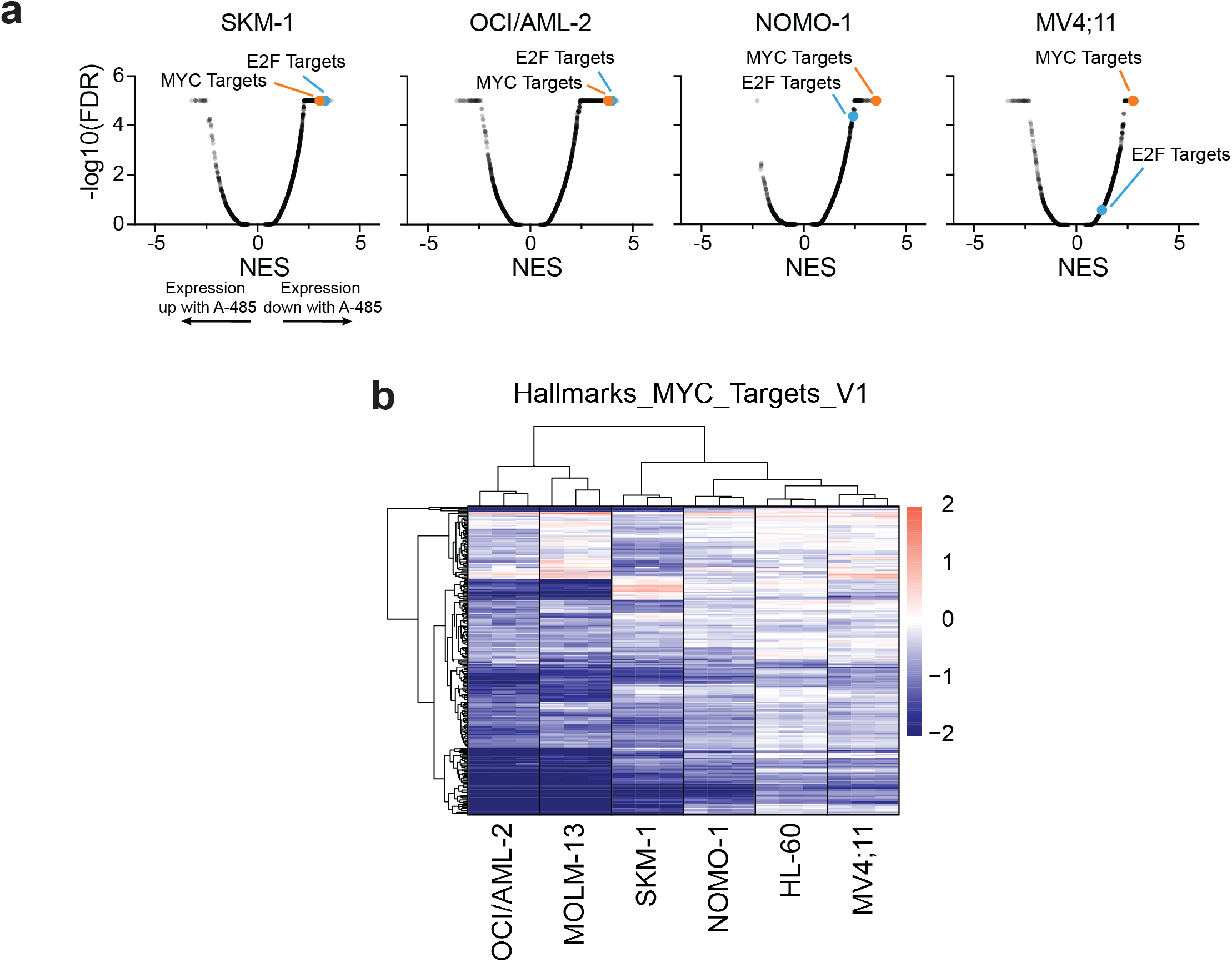
Secondary transcriptional responses to CBP/p300 HAT inhibition in a panel of AML cell lines. **a**, Gene set enrichment analysis for changes in gene expression following 48-h of A-485 (3 μM) treatment. **b**, Heatmap representation of DMSO-normalized fold changes in spike-in normalized gene expression of MYC target genes caused by treatment with A-485 (3 μM).

**Extended Data Figure 5:**
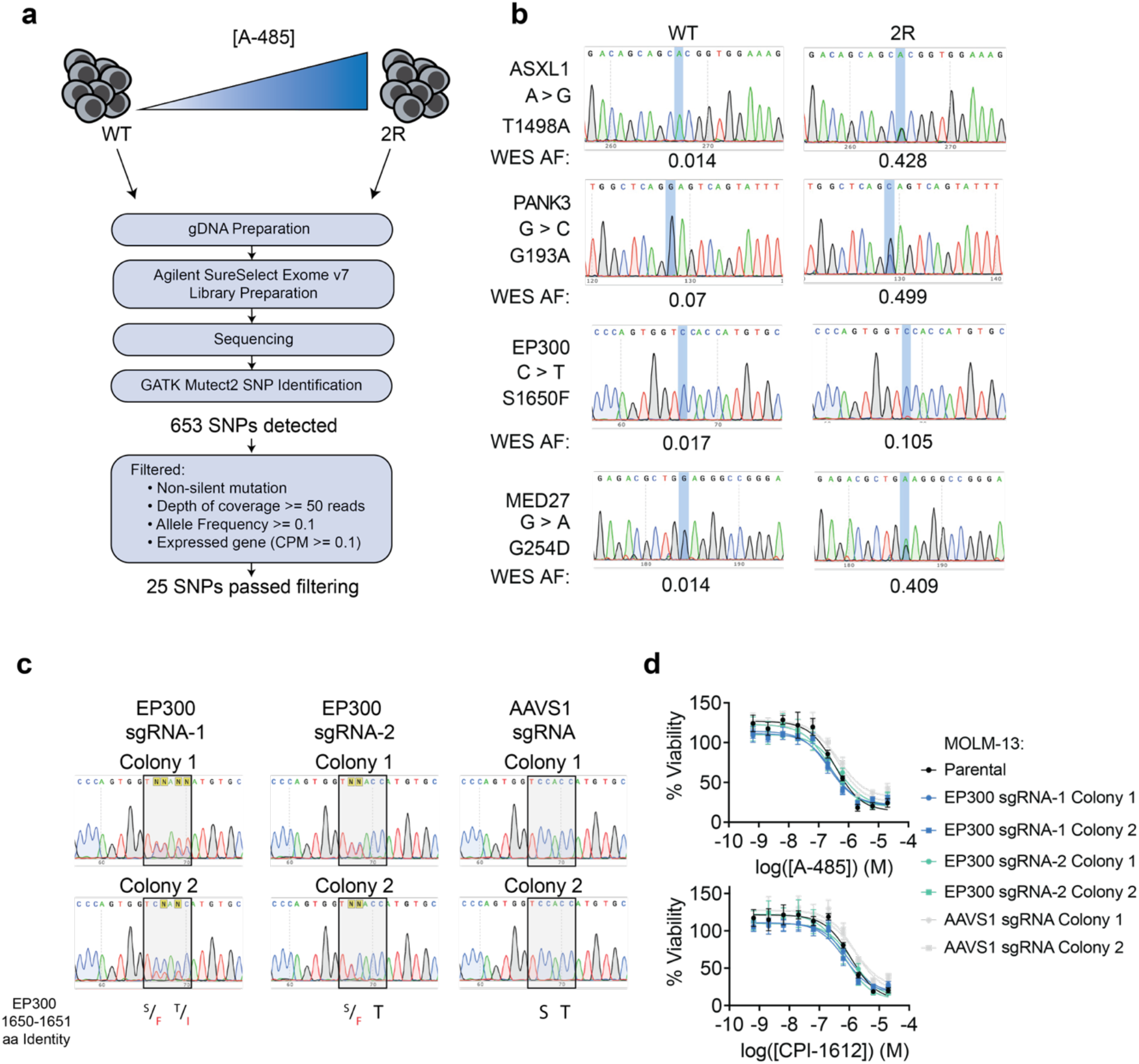
Whole-exome sequencing to identify variantss that contribute to HAT inhibitor resistance. **a**, Schematic of whole-exome sequencing experimental and data analysis pipeline. **b**, Sanger sequencing traces confirming SNPs identified in MOLM-13-2R through WES. Indicated allele frequencies were predicted from WES. **c**, Sanger sequencing traces confirming base-editing activity at codon for EP300 S1650. **d**, DMSO-normalized cellular viability in MOLM-13 expressing CRISPR-directed base editor and EP300-targeting sgRNA or control AAVS1-targeting sgRNA. Mean ± SD, n = 3.

**Extended Data Figure 6:**
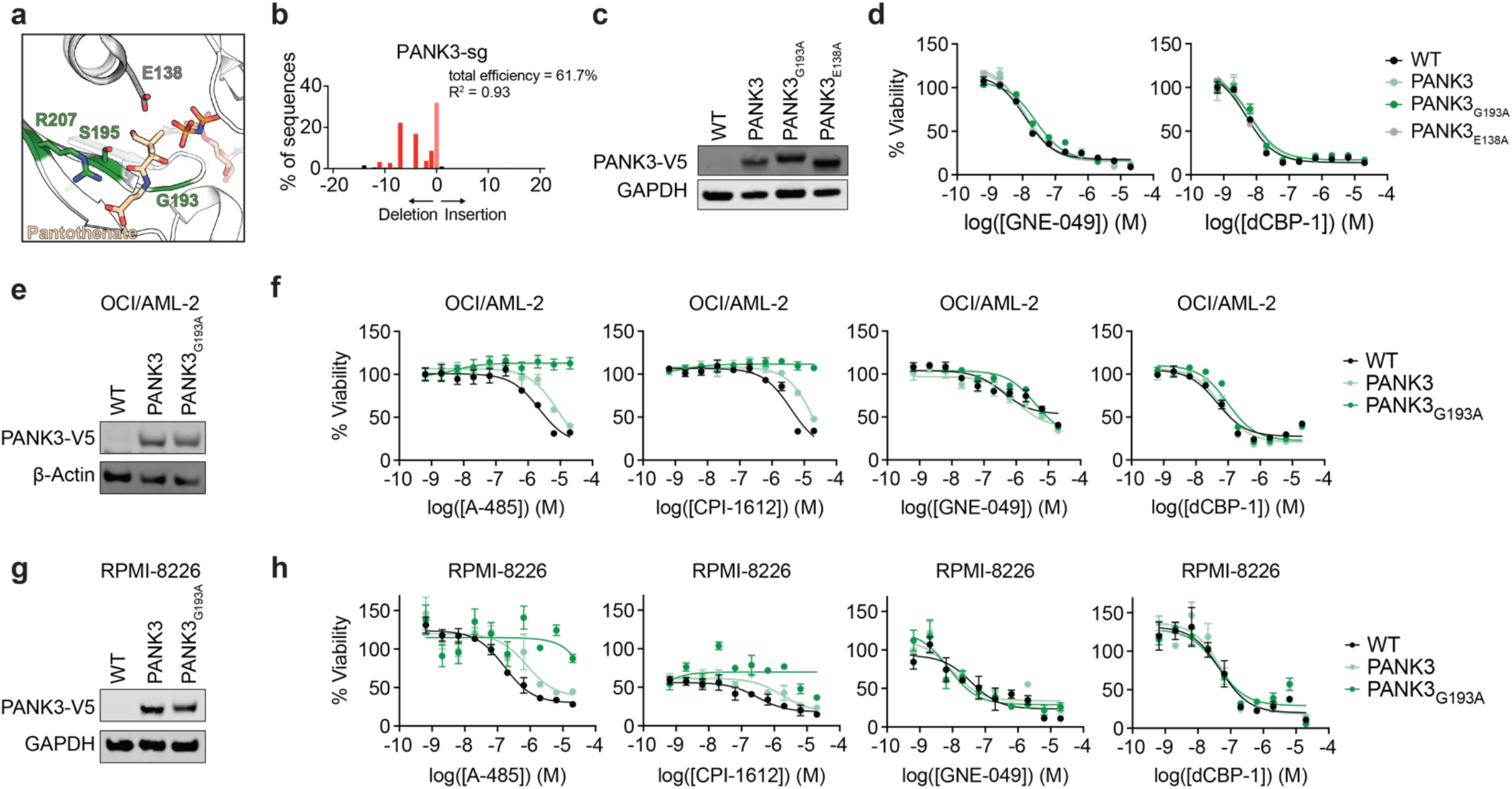
PANK3 and PANK3_G193A_ expression confer resistance to CBP/p300 HAT inhibitors. **a**, Structural view of the acetyl-CoA and pantothenate binding site in PANK3 (PDB: 5KPR). Amino acids G193, S195V, and R207 are highlighted in green. **b**, Validation of sgRNA targeting PANK3 by TIDE 4-d post-transduction. **c**, Immunoblot confirmation of overexpression of PANK3, PANK3_G193A_, and PANK3_E183A_ in MOLM-13. **d**, DMSO-normalized cellular viability in MOLM-13 expressing PANK3 ORFs after 72-h treatment with indicated compounds as determined by ATP-lite assay. Mean ± SEM, n = 3. **e**, Immunoblot confirmation of overexpression of PANK3 and PANK3_G193A_ in OCI/AML-2. **f**, DMSO-normalized cellular viability in OCI/AML-2 expressing PANK3 ORFs after 72-h treatment with indicated compounds as determined by ATP-lite assay. Mean ± SEM, n = 3. **g**, Immunoblot confirmation of overexpression of PANK3 and PANK3_G193A_ in RPMI-8226. **h**, DMSO-normalized cellular viability in RPMI-8226 expressing PANK3 ORFs after 72-h treatment with indicated compounds as determined by ATP-lite assay. Mean ± SEM, n = 3.

**Extended Data Figure 7:**
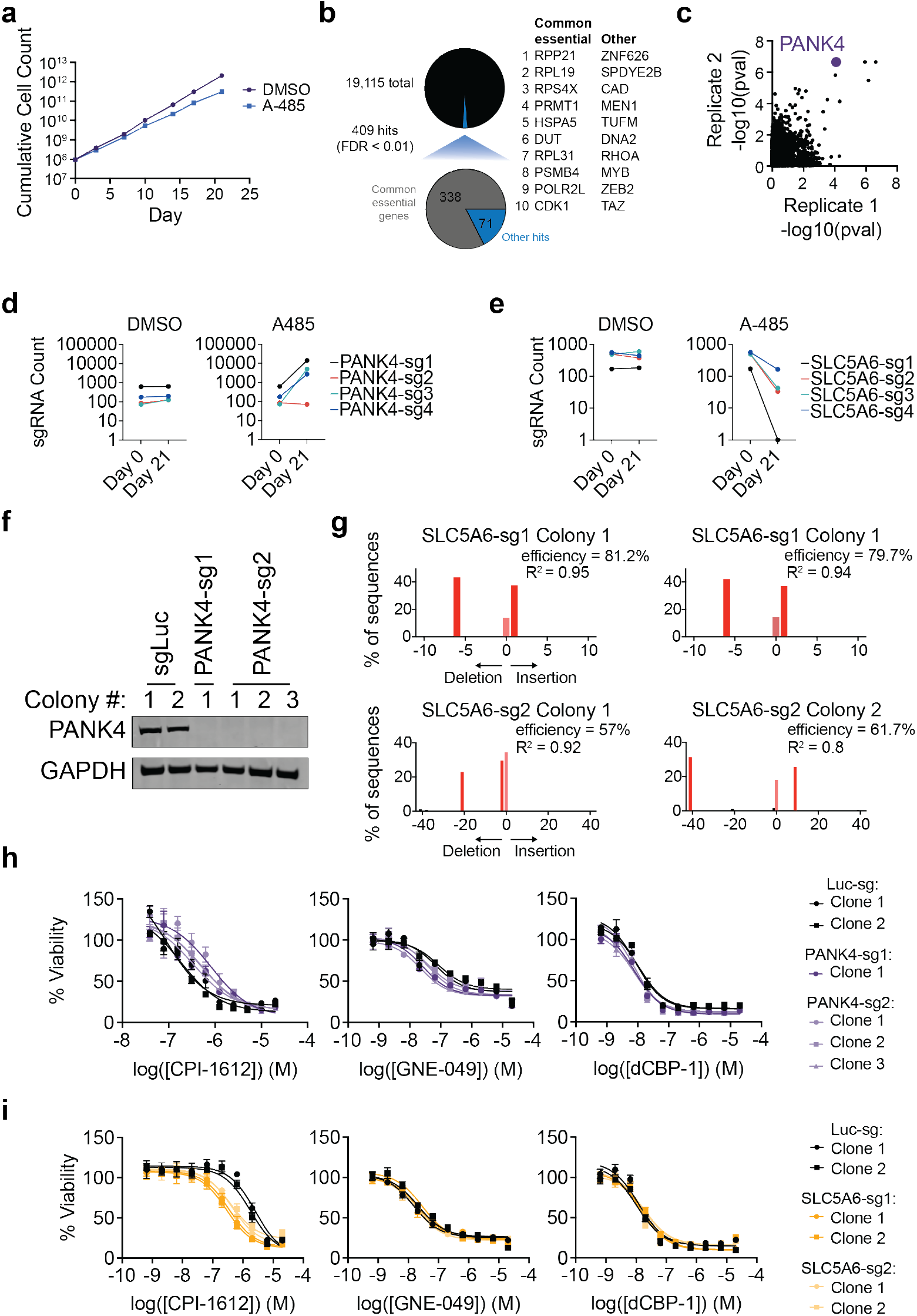
Genome-wide CRISPR screen identifies genes in CoA biosynthesis pathway as modulators of HAT inhibitor sensitivity. **a**, Growth curves for genome-wide CRISPR-Cas9 screen. **b**, Summary of dropout in CRISPR-Cas9 knock-out screen after 21-d of culture in DMSO showing the number of hits (FDR < 0.01) and the percentage of hits known to be common essential genes. **c**, Enrichment of genes from two replicate CRISPR-Cas9 screens in MM1.S treated with A-485. Data accessed from GSE173772. **d**, Counts from sgRNAs targeting *PANK4* from CRISPR-Cas9 modifier screen in MOLM-13. **e**, Counts from sgRNAs targeting *SLC5A6* from CRISPR-Cas9 modifier screen in MOLM-13. **f**, Immunoblot confirmation of PANK4 knockout in clonal populations expressing PANK4-targeting sgRNAs in MOLM-13. **g**, Confirmation of SLC5A6 gene locus disruption in clonal populations expressing SLC5A6-targeting sgRNAs in MOLM-13 by TIDE. **h**, DMSO-normalized cellular viability in MOLM-13 clonal populations expressing PANK4-targeting sgRNAs after 72-h treatment with indicated compounds as determined by ATP-lite assay. Mean ± SEM, n = 3. **i**, As in **f**, but in MOLM-13 clonal populations expressing SLC5A6-targeting sgRNAs. Mean ± SEM, n = 3.

**Extended Data Figure 8:**
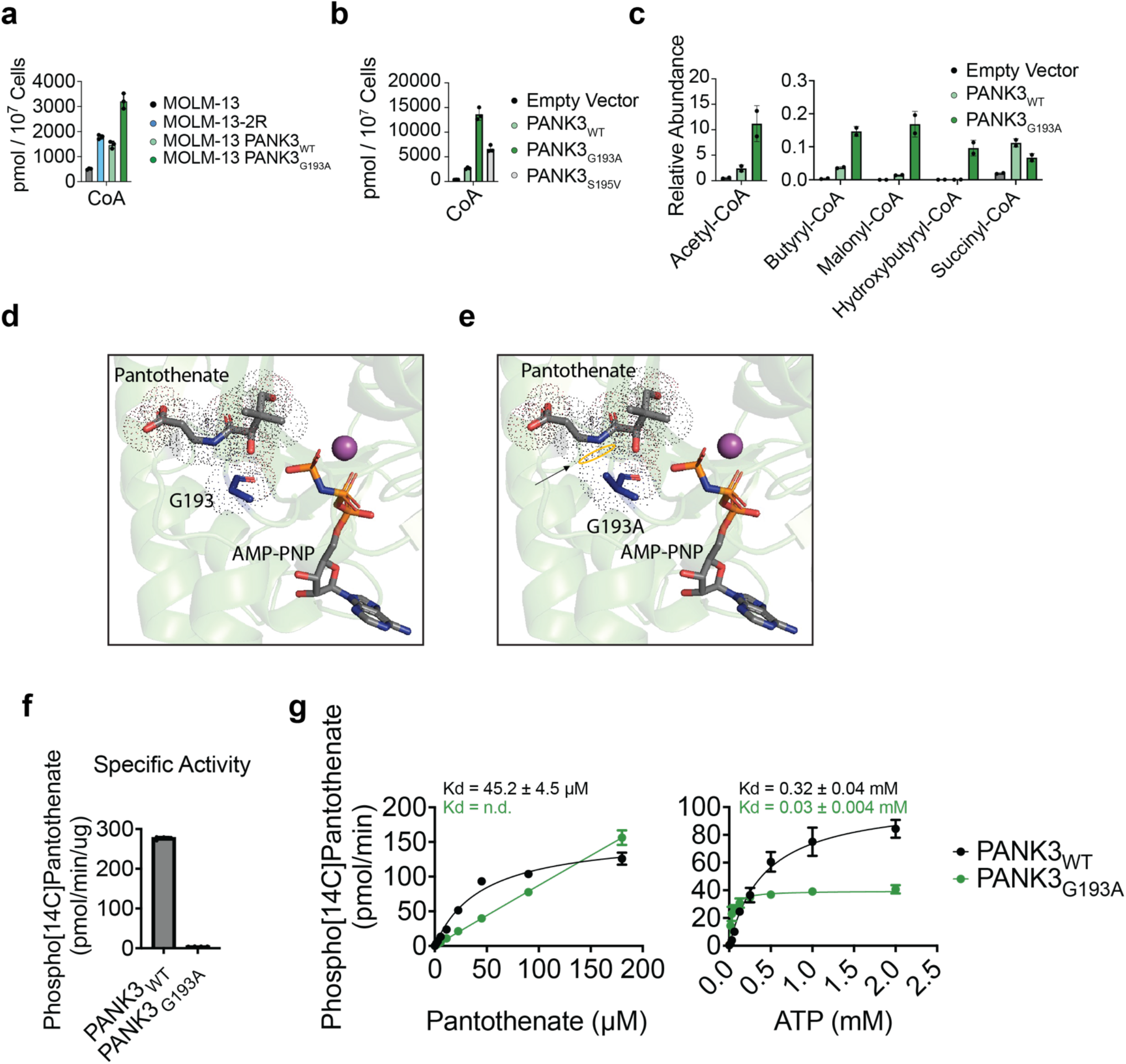
PANK3_G193A_ is refractory to feedback inhibition from acetyl-CoA. **a**, Total cellular CoA levels determined by mass-spectrometry. Mean ± SD, n = 3. **b**, Total cellular CoA levels determined by mass-spectrometry in HEK293T cells expressing PANK3 ORFs. Mean ± SD, n = 3. **c**, Relative levels of acyl-CoA species determined by mass spectrometry in HEK293T cells expressing PANK3 ORFs. Mean ± SD, n = 2. **d**, Structural view of pantothenate and ATP binding site in PANK3 (PDB: 5KPR). **e**, Structural view of pantothenate and ATP binding site in PANK3 with modeling of the G193A mutation. Predicted steric clash indicated by arrow and yellow circle. **f**, Specific activity of purified PANK3 and PANK3_G193A_ in radiometric PANK activity assay. Mean ± SD, n = 4. **g**, Kinetic analysis of PANK3 and PANK3_G193A_ activity with respect to pantothenate (left) and ATP (right) concentrations. Mean ± SD, n = 4.

**Extended Data Figure 9:**
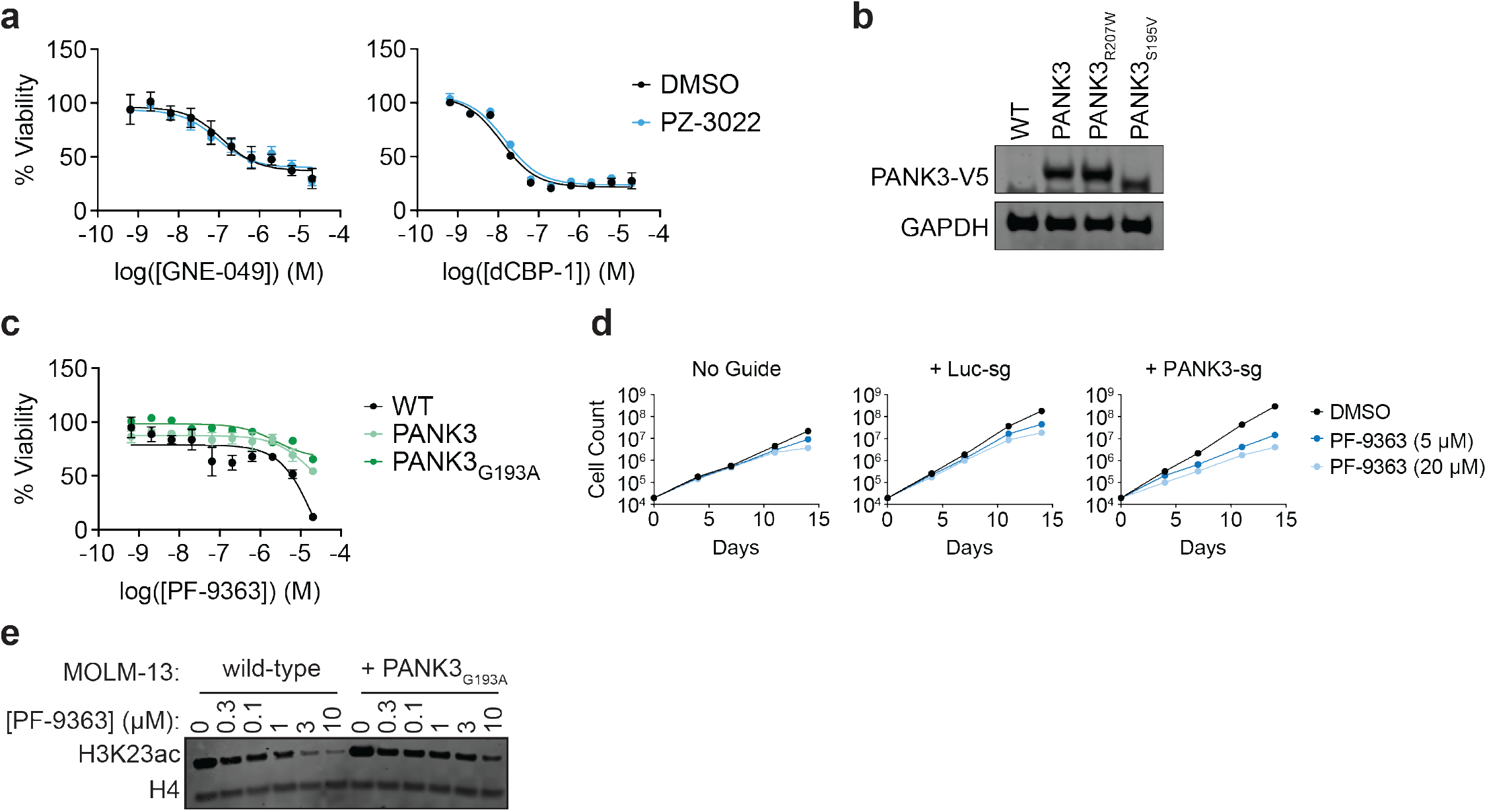
PANK3 hyperactivity confers resistance broadly to HAT inhibitors. **a,** DMSO-normalized cellular viability in MOLM-13 co-treated with indicated compound and either PZ-3022 (1 μM) or DMSO for 72-h. Mean ± SEM, n = 3. **b**, Immunoblot confirmation of overexpression of PANK3, PANK3_R207W_, and PANK3_S195V_ in MOLM-13. **c**, DMSO-normalized cellular viability in MOLM-13 expressing PANK3 ORFs after 72h treatment with PF-9363 as determined by ATP-lite assay. **d**, Proliferation of MOLM-13-2R expressing indicated sgRNAs over 14-d of PF-9363 treatment. **e**, Immunoblot for KAT6A/B substrate H3K23ac following 6 hours of PF-9363 treatment.

